# Kinetics of cytokine receptor trafficking determine signaling and functional selectivity

**DOI:** 10.1101/638031

**Authors:** J. Martinez-Fabregas, S. Wilmes, L. Wang, M. Hafer, E. Pohler, J. Lokau, C. Garbers, A. Cozzani, J. Piehler, M. Kazemian, S. Mitra, I. Moraga

## Abstract

Cytokines activate downstream signaling networks via assembly of cell surface receptors, but it is unclear whether modulation of cytokine-receptor binding parameters can modify biological outcomes. We have engineered variants of IL-6 with different affinities to the gp130 receptor chain to investigate how cytokine receptor binding kinetics influence functional selectivity. Engineered IL-6 variants showed a range of signaling amplitudes, from minimal to full agonist, and induced biased signaling, with changes in receptor binding kinetics affecting more profoundly STAT1 than STAT3 phosphorylation. We show that this differential signaling arises from defective translocation of ligand-gp130 complexes to the endosomal compartment and competitive STAT1/STAT3 binding to phospho-tyrosines in gp130, and results in unique patterns of STAT3 binding to chromatin. This, in turn, leads to a graded gene expression response and substantial differences in *ex vivo* differentiation of Th17, Th1 and Treg cells. These results provide a molecular understanding of signaling biased by cytokine receptors, and demonstrate that manipulation of signaling thresholds is a useful strategy to decouple cytokine functional pleiotropy.

## INTRODUCTION

Cytokines modulate the immune response by activating a common JAK/STAT signaling pathway upon cell surface receptor dimerization/oligomerization (Gorby et al., 2018; Stroud and Wells, 2004; Wang et al., 2009; Wells et al., 1993). A conundrum in the field pertains to how biological specificity is achieved in the cytokine system by using such reduced number of signaling intermediaries, i.e. four JAKs and seven STATs (Murray, 2007; Schindler et al., 2007). Indeed, there are numerous examples in the literature where cytokines activating the same STATs in CD4 T cells, e.g. IL-6 and IL-10 (Grötzinger et al., 1997; Walter, 2004), produce opposite responses, i.e. pro-inflammatory vs anti-inflammatory responses respectively (Hunter and Jones, 2015; Wilson et al., 2005).

In recent years, a number of studies in multiple cytokine systems have shown that cytokine signaling is not an “all or none” phenomenon and can be modulated by alterations of cytokine-receptor binding properties (Spangler et al., 2015). Changes in cytokine-receptor binding kinetics and strength were shown to play a crucial role in defining type I and type III interferons biological potencies (Mendoza et al., 2017; Pestka, 2007; Piehler et al., 2012; Subramaniam et al., 1995). A mutation in erythropoietin (Epo) found in humans, which reduces its binding affinity for its receptor (EpoR), was shown to biased signaling output by EpoR and caused severe anemia in human patients (Kim et al., 2017). Biased EpoR signaling was also achieved using surrogate Epo ligands that altered the receptor binding topology (Moraga et al., 2015b). Cross-reactive cytokine-receptor systems, where shared receptors engage multiple cytokines and elicit differential responses is another example where receptor binding properties influence signaling and activity, e.g. the IL-4/IL-13 system (Heller et al., 2008; LaPorte et al., 2008), IL-2/IL-15 system (Ring et al., 2012; Rochman et al., 2009; Waldmann, 2006) and the IL-6 family system (Wang et al., 2009). Viruses often encode for cytokine-like proteins that bind cytokine receptors with altered binding properties, providing them with means to fine-tune the immune response to their own advantage (Boulanger et al., 2004; Walter, 2004). All these examples strongly argue in favour of cytokine-receptor binding parameters contributing to regulate signaling, however a model providing molecular bases for signaling biased by cytokine receptor is missing.

Biased signaling is not a unique feature of the cytokine family. G-protein-coupled receptors (GPCRs), which contain seven-transmembrane domains, are the classical system where biased signaling was first described (Hilger et al., 2018; Wootten et al., 2018). In this system, different ligands binding a common receptor can trigger differential signaling programs by instructing specifics allosteric changes in the transmembrane α-helices of the receptor (Hilger et al., 2018; Wootten et al., 2018). However, although elegant, this mechanism cannot be applied to the family of single-spanning transmembrane domain cytokine receptors. How can cytokine receptors trigger biased signaling responses then? A common feature to all cytokine systems is that upon ligand stimulation cytokine-receptor complexes traffic to intracellular compartments, where they are often degraded, contributing to switching off the response (Becker et al., 2010; Bulut et al., 2011; Claudinon et al., 2007; German et al., 2011; Keeler et al., 2007; Shah et al., 2006). However, a complex positive regulatory role of endocytosis in cytokine signaling has emerged (Becker et al., 2010; Cendrowski et al., 2016; Fallon and Lauffenburger, 2000; Marchetti et al., 2006; Sarkar et al., 2002). Several studies recently suggested a novel role for the endosomal compartment in stabilizing cytokine receptor dimers by enhancing local receptor concentrations (Gandhi et al., 2014; Moraga et al., 2015a), thus contributing to signaling fitness even at low complex stabilities. In agreement with this model, mutations on cytokine receptors that alter their intracellular traffic can result in activation of novel or deregulated signaling programs causing disease (Reddy et al., 1996). Furthermore, activated JAK/STAT proteins have been described in endosomes after interferon stimulation, suggesting that signaling continues upon receptor internalization (Payelle-Brogard and Pellegrini, 2010). How changes in cytokine-receptor complex half-life and endosomal trafficking fine-tunes cytokine signaling and biological responses requires further investigation.

Here, using model cell lines and primary human CD4 T cells, we systematically explored how modulation of cytokine-receptor complex stability impacts signaling identity and biological responses, using IL-6 as a model system. IL-6 is a highly pleiotropic cytokine, which critically contributes to mounting the inflammatory response (Grötzinger et al., 1997; Hunter and Jones, 2015; Naka et al., 2002). IL-6 stimulation drives differentiation of Th-17 cells (Jones et al., 2010; Kimura and Kishimoto, 2010; Louten et al., 2009), and inhibits the differentiation of Th-1 (Diehl and Rincon, 2002) and T regulatory (reg) cells (Kimura and Kishimoto, 2010; Korn et al., 2008). Deregulation of IL-6 levels and activities is often found in human diseases, making IL-6 a very attractive therapeutic target (Hunter and Jones, 2015). IL-6 exerts its immune-modulatory activities by engaging a hexameric complex comprised of two molecules of IL-6Rα, two molecules of gp130 and two molecules of IL-6, leading to the downstream activation of STAT1 and STAT3 transcription factors (Wang et al., 2009). Using the yeast-surface display engineering platform, we isolated a series of IL-6 variants binding gp130 with different affinities, ranging from wild-type binding affinity to more than 2000-fold enhanced binding. Quantitative signaling and imaging studies revealed that reduction in cytokine-receptor complex stability resulted in differential cytokine-receptor complex dynamics, which ultimately led to activation of biased signaling programs. Low affinity IL-6 variants, failed to translocate to intracellular compartments and induce gp130 degradation, triggering STAT3 biased responses. Indeed, inhibition of gp130 intracellular translocation by chemical or genetic blockage of clathrin-mediated traffic, reduced STAT1 activation levels, without affecting STAT3 activation. Through a series of molecular and cellular assays we demonstrated that STAT1 requires a higher number of phospho-Tyr available in gp130 to reach maximal activation, explaining its enhanced sensitivity to changes in cytokine-receptor complex stability. The biased signaling programs engaged by the IL-6 variants did not have a linear effect on STAT3 transcriptional activities. Reduced STAT3 activation levels by the low affinity IL-6 variants resulted in graded STAT3 binding to chromatin and gene expression, with some genes exhibiting a high degree of sensitivity to STAT3 activation levels, and other genes being equally induced by all three IL-6 variants. Moreover, IL-6 immuno-modulatory activities exhibited different sensitivity thresholds to changes on STAT activation levels, with Th-17 differentiation being induced by all three variants, and inhibition of Treg and Th-1 differentiation only robustly promoted by the high affinity variant. Our results provide a molecular model using spatio-temporal dynamics of cytokine-receptor complexes and competitive binding of STATs proteins for phosphorylated tyrosine residues, to explain how cells integrate cytokine signaling signatures into specific biological responses through the establishment of different gene induction thresholds. At the more practical level, our results highlight that manipulation of cytokine-receptor binding parameters via protein engineering is a useful strategy to decouple cytokine functional pleiotropy, a major source of toxicity in cytokine-based therapies.

## RESULTS

### Engineering IL-6 variants with different binding affinities for gp130

IL-6 stimulates signaling via a hexameric receptor comprised of two copies of IL-6, two gp130 and two IL-6Rα receptor subunits (Wang et al., 2009). To understand whether variations on cytokine-receptor binding parameters would instruct different biological outcomes, we used yeast surface display to engineer a series of IL-6 variants binding gp130 with different affinities. IL-6 interacts with gp130 on two sites, named site-2, which uses helixes A and C on IL-6, and site-3 which uses part of the AB-loop and helix D (Fig. 1 a) (Wang et al., 2009). We focused on site-2 since this site is the main driver of gp130-IL6 interaction. Using the existing crystal structure of the IL-6 hexameric complex, we identified 14 amino acids on IL-6 forming the site-2 binding interface, which we randomized using a ‘NDT’ degenerate codon encoding amino acids: G,V,L,I,C,S,R,H,D,N,F,Y (Fig. 1 a). The resulting library contained more than 3×10^8^ unique variants.

**Figure 1:**
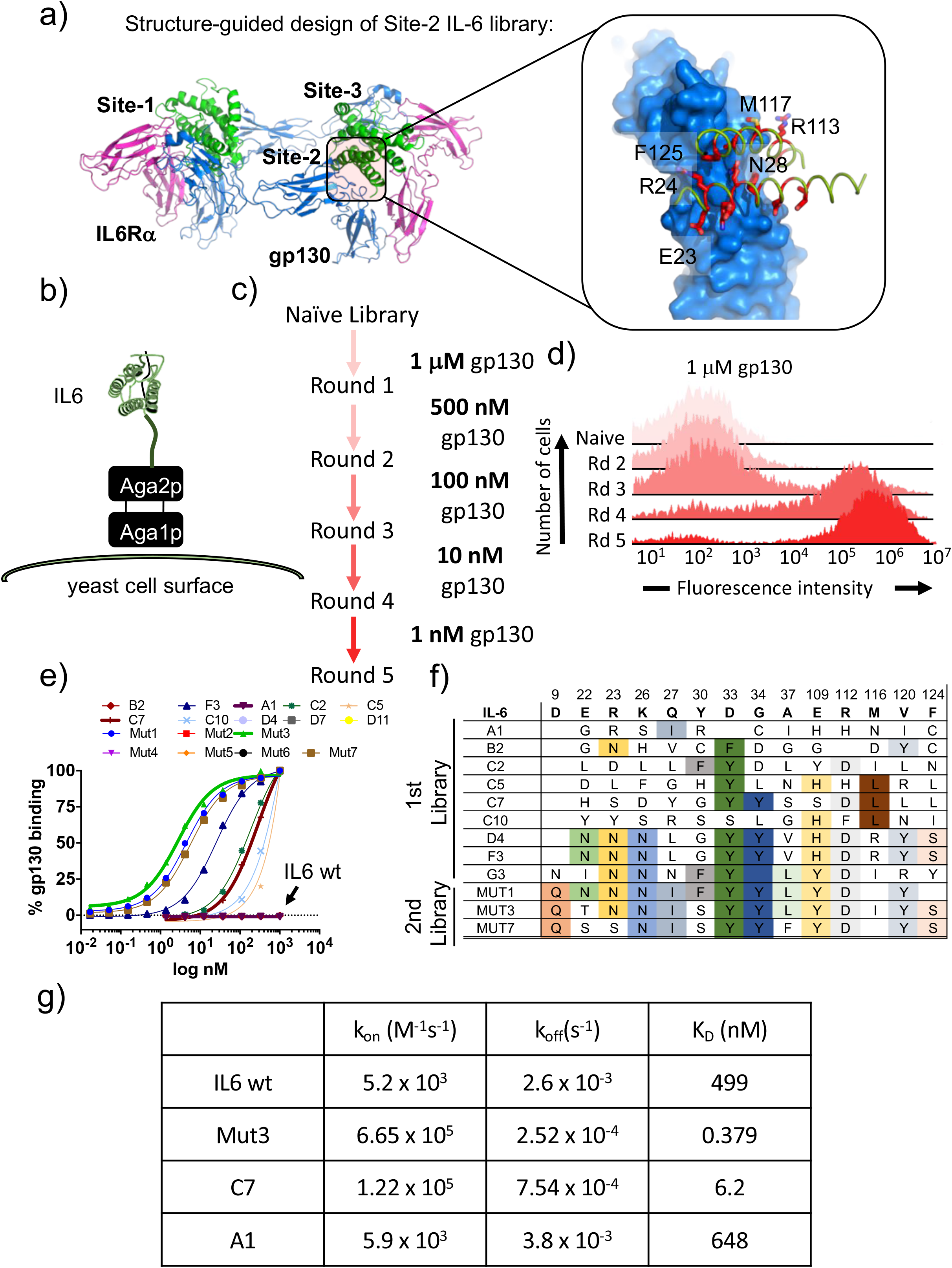
Isolation of IL-6 variants binding gp130 with different affinities. (a) Crystal structure of IL-6, in green, bound to gp130 and IL-6Rα ectodomains, in blue and pink respectively. Inlet highlights the IL-6/gp130 site-2 binding interface. Amino acids included in the library design are colored in red. (b) Schematic representation of IL-6 display in the yeast surface via aga2p-aga1p interaction. (c) Work-flow of IL-6 library selection process. Five rounds of selection were undertaken, starting with 1 μM of gp130 ectodomain and finishing with 1 nM. (d) Representative gp130 staining of the selected IL-6 library. The five rounds of selections were incubated with 1 μM of biotinylated gp130 for 1 hr followed by 15 min incubation with SA-alexa647. Early rounds exhibit weak binding to gp130, but as the library converged into few high affinity clones, the gp130 staining improve significantly. (e) Dose/Response gp130 binding curves performed in single yeast colonies, each encoding a different IL-6 variant. Gp130 concentration started at 1 μM, and eight different concentrations in a 1/3 dilution series were tested. (f) Amino acid sequences corresponding to isolated IL-6 variants. Variants from early rounds are displayed on the top of the table and exhibit a wider range of mutations. As the library converges, fewer unique sequences are found, with all of them exhibiting similar mutations as at the bottom of the table. (g) Table illustrating K_on_, K_off_ and K_D_ binding constants obtained from surface plasmon resonance studies for IL-6 wt and Mut3, C7 and A1 IL-6 variants.

The library was selected for gp130 binders through five rounds of selection in which the gp130 concentration was gradually decreased from 1 μM to 1 nM (Fig 1 b-d). Nine clones were selected based on their on-yeast binding titrations and their sequences were obtained (Fig 1 e-f). From this initial library, IL-6 variants exhibiting a wide range of binding affinities for gp130 were isolated, ranging from wt affinity (A1) to 200-fold better binding (F3) (Fig. 1 e). In order to isolate IL-6 variants binding with even higher affinity to gp130, we performed a second library, where we further engineered the F3 mutant, by carrying out a soft randomization of the amino acids forming the gp130 site-2 binding interface. After five additional rounds of selection we isolated three new variants (Mut1, Mut3 and Mut7) that bound gp130 with an apparent binding K_D_ of 2 nM (Fig. 1 e-f). The binding affinities of three of those variants, belonging to the first and second libraries, were confirmed by surface plasmon resonance (SPR) studies, with values ranging from 648 nM (A1) to 6.2 nM (C7) and 379 pM (Mut3) (Fig 1 g and Sup. Fig 1-d).

### IL-6 variants induce differential STAT3/STAT1 activation ratios

IL-6 binding to gp130 triggers the phosphorylation and activation of STAT1 and STAT3 factors via JAK1. Next we studied the different STAT1 and STAT3 activation signatures elicited by the A1, C7 and Mut3 IL-6 variants. For that, we used HeLa cells, which express very low levels of IL-6Rα subunit and therefore allow us to study the contribution of gp130 binding to signaling output by the IL-6 variants. As control we used IL-6 wild type (wt), which requires IL-6Rα expression to activate signaling, and Hyper IL-6 (HyIL-6), which binds gp130 with high affinity and potently triggers signaling in cells lacking IL-6Ra (Fischer et al., 1997). The three engineered variants exhibited different degrees of IL-6Rα dependency based on their gp130 binding affinities (Sup. Fig. 1 e-f). As expected, while IL-6 wt stimulation led to a poor signaling response in HeLa cells, HyIL-6 stimulation produced a robust STAT1 and STAT3 activation in dose-response studies (Fig. 2 a-b). Interestingly, different IL-6 variants drove differential phosphorylation amplitudes in STAT1 and STAT3 (Fig. 2 a-b). These differences in signaling amplitudes could not be rescued by further increases in ligand concentration (Fig. 2 a-b), nor were the result of altered signaling kinetics induced by the IL-6 variants (Fig. 2 c-d). Strikingly, we observed that STAT1 phosphorylation was profoundly more affected than STAT3 by changes in gp130 binding affinities (Fig. 2 a-d). While Mut3 activated STAT1 and STAT3 to the same extent than HyIL-6, the C7 variant induced 70% of the STAT3 phosphorylation levels but only 25% of the STAT1 phosphorylation levels induced by HyIL-6. Similarly, the A1 variant induced 50% of the STAT3 phosphorylation levels as compared to HyIL-6, but failed to induce STAT1 phosphorylation (Fig.2 a-d). This biased STAT3 activation by the IL-6 variants resulted in altered STAT3/STAT1 activation ratios, with IL-6 variants binding with lower affinity to gp130 exhibiting a disproportionally high activation of STAT3 versus STAT1 (Fig 2 e and Sup. Fig. 1g).

**Figure 2:**
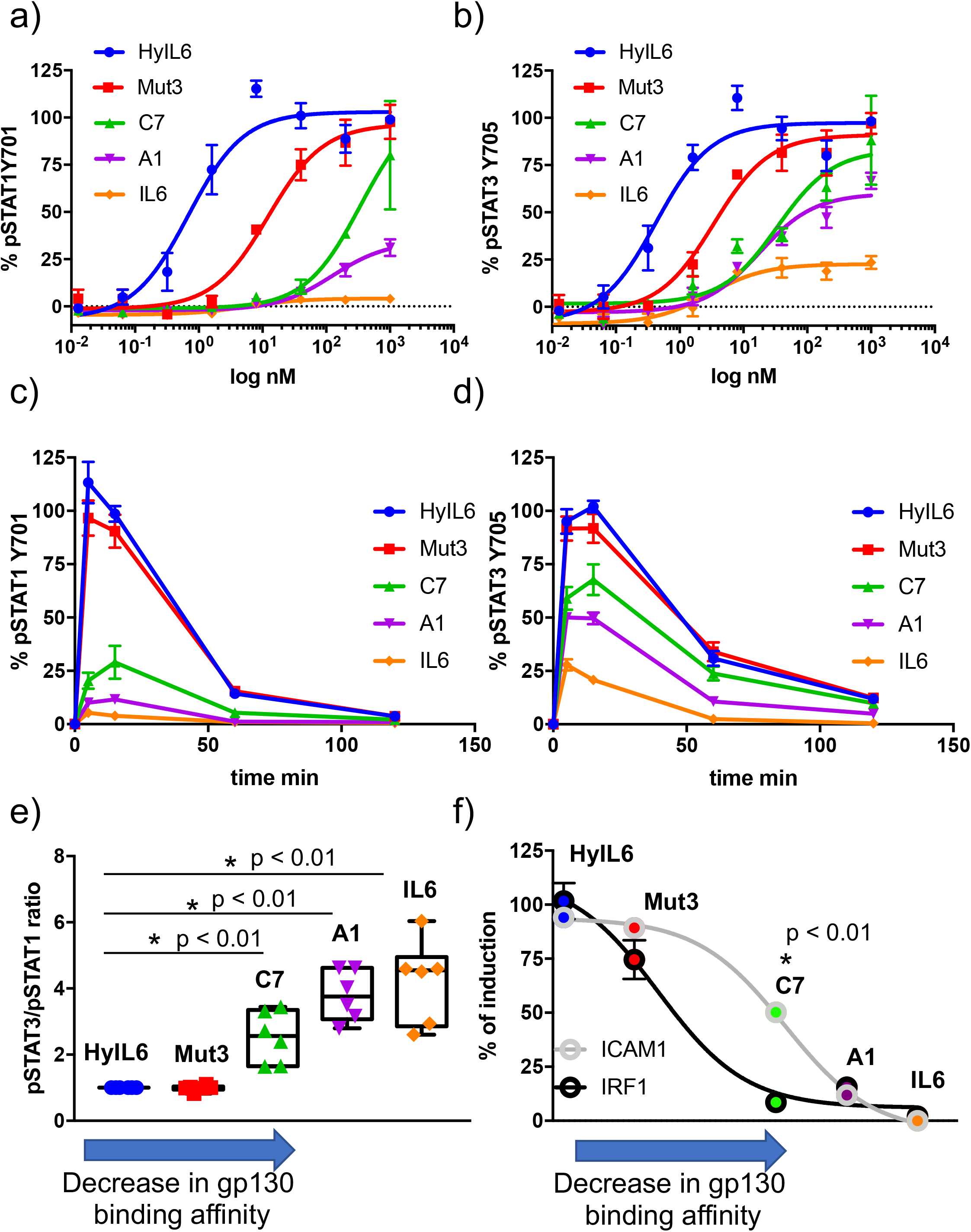
Determination of signaling signatures activated by IL-6 variants. (a-b) HeLa cells were stimulated with the indicated doses of IL-6 ligands for 15 min and levels of STAT1 (a) and STAT3 (b) were analyzed by phospho-Flow cytometry. Sigmoidal curves were fitted with GraphPath Prism software. Data are mean +/− SEM from three independent replicates, each performed in duplicate. (c-d) HeLa cells were stimulated with 100 nM of IL-6 ligands for the indicated times and the levels of STAT1 (c) and STAT3 (d) were analyzed by phospho-Flow cytometry. Data are mean +/− SEM from three independent replicates, each performed in duplicate. (e) Differential STAT activation by engineered IL-6 ligands. pSTAT3/pSTAT1 ratios are plotted for all the IL-6 ligands. An arrow indicating the binding affinity trends of each ligand was placed in the X axis of the plot. Low gp130 affinity ligands exhibit a more pronounced STAT3/STAT1 ratio than high affinity ligands. Data are mean +/− SEM from three independent replicates, each performed in duplicate. (f) Comparison of STAT1- (IRF1) and STAT3-dependent (ICAM1) gene induction by engineered IL-6 ligands. HeLa cells were stimulated with saturating concentrations (100 nM) of the different IL-6 ligands for either 2 hrs (IRF1) or 24 hrs (ICAM1) and the levels of IRF1 and ICAM1 induction were measured via flow cytometry. Data are mean +/− SEM from three independent replicates, each performed in duplicate. An arrow indicating the binding affinity trends of each ligand was placed in the X axis of the plot.

To investigate whether the biased STAT3 signature induced by the different IL-6 variants would impact their transcriptional programs, we analysed the induction of a classical STAT1-dependent and STAT3-dependent proteins, i.e. IRF1 and ICAM-1, by the three IL-6 variants (Gil et al., 2001; Wung et al., 2005). HeLa cells were stimulated with saturating concentrations of the different variants and the levels of IRF1 and ICAM1 expression were measured by flow cytometry. As shown in Fig. 2 f, induction of IRF1 expression was more sensitive to changes on gp130 binding affinity, paralleling the sensitivity of STAT1 activation. Overall these data indicate that modulation of cytokine-receptor binding parameters decouples signaling output and transcriptional programs.

### Short-lived IL-6-gp130 complexes fail to traffic to intracellular compartments

We have shown that engineering IL-6 to display different affinities for gp130 results in biased STAT3 and STAT1 responses by this receptor system. However, the molecular basis that allow a single-pass transmembrane receptor, such as gp130, to fine tune its signaling output in response to changes in binding energy remains unclear. We reasoned that one of the first steps affected by changes in gp130 binding affinity would be the assembly kinetics of the IL-6/gp130 hexameric complex in the cell membrane. To study this step, we probe the assembly of gp130 dimers at the single molecule level using total internal reflection fluorescence (TIRF) microscopy. In these experiments, we transfected HEK293 cells lacking endogenous gp130 expression (Schwerd et al., 2017) with gp130 N-terminally tagged with a meGFP, which was rendered non-fluorescent by the Y67F mutation (Fig. 3 a). This tag (mXFP) is recognized by dye-conjugated anti-GFP nanobodies, allowing quantitative fluorescence labelling of gp130 at the cell surface of live cells. Well-balanced dual-colour labelling was achieved using equal concentrations of nanobodies either conjugated with RHO11 or with DY647 as recently shown in (Kim et al., 2017; Moraga et al., 2015b). Diffusion and interaction of individual gp130 in the plasma membrane was probed by dual-colour total internal refection fluorescence (TIRF) microscopy. Single-molecule co-localization and co-tracking analysis was used to identify correlated motion (co-locomotion) of the spectrally separable fluorescent molecules, which was taken as a readout of productive dimerization of gp130. Substantial gp130 dimerization could only be discerned after addition of HyIL-6 (Fig. 3 b). Under these conditions, co-trajectories corresponding to individual gp130 dimers co-locomoting for more than 10 consecutive frames (~320ms) were observed (Sup. Fig.2 a-b). We detected significant ligand-induced gp130 co-locomotion for all IL-6 variants except for A1, with levels of co-locomotion paralleling gp130 binding affinities, i.e. HyIL-6>Mut3>C7>A1 (Fig. 3 b). In agreement with this, we observed a strong decrease in lateral diffusion mobility, which can be ascribed to receptor dimerization (Moraga et al., 2015b; Wilmes et al., 2015) (Fig.3 c).

**Figure 3:**
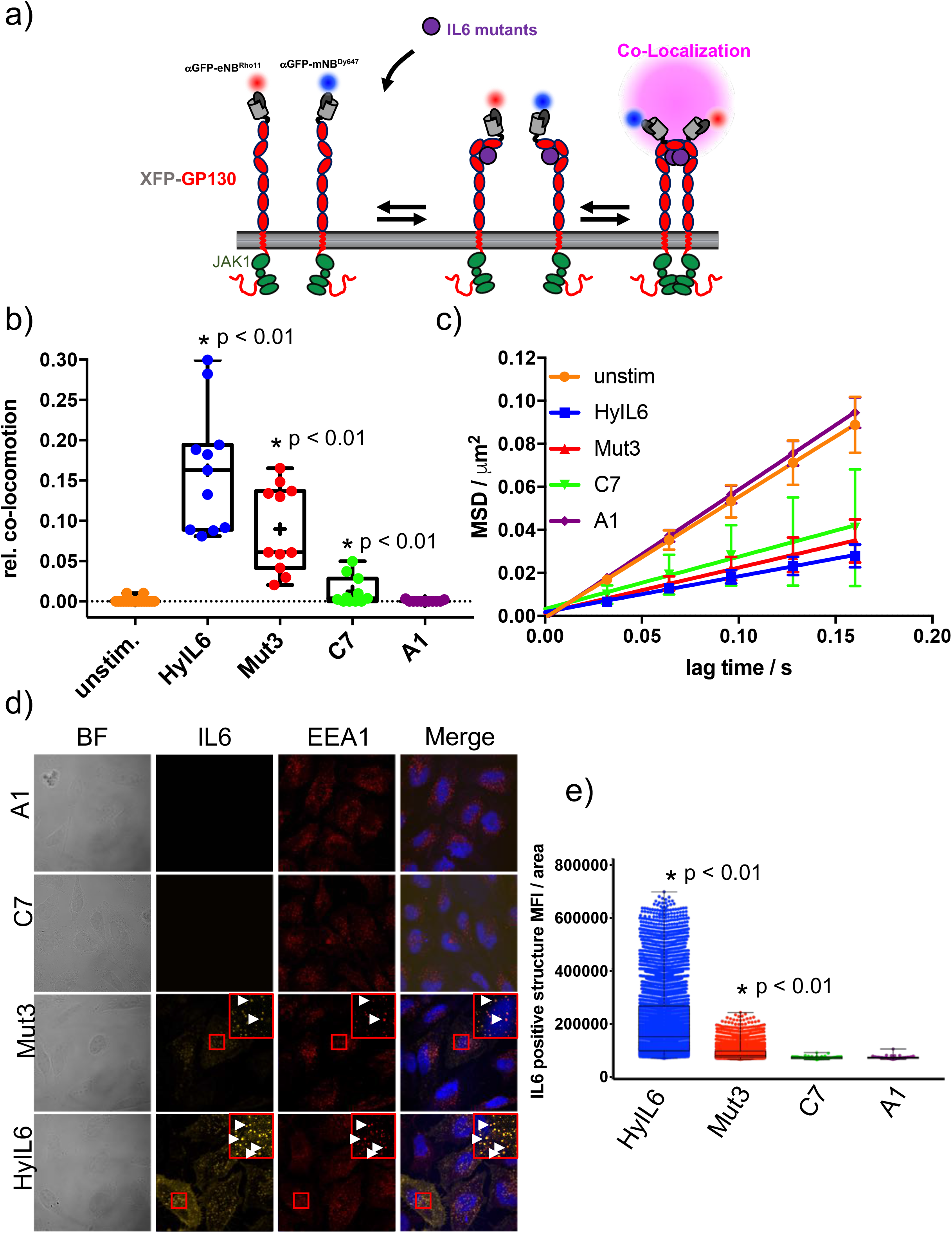
gp130 cell surface dynamics induced by the different IL-6 variants. (a) Quantification of gp130 homodimerization in the plasma membrane by dual-color singlemolecule co-localization/co-tracking. mXFP-gp130 was expressed in HEK 293T gp130 k/o cells and labeled via anti-GFP nanobodies conjugated with RHO11 and DY647, respectively. (b) Relative amount of co-trajectories for unstimulated gp130 and after stimulation with HY-IL6 and IL-6 mutants (Mut3, C7 and A1). (c) Mean square displacement (MSD) analysis of mXFPm-gp130 diffusion properties in absence of ligand and in presence of HyIL-6, or IL-6 mutants respectively (1-5 steps). (d) Uptake of DY547-conjugated HyIL-6 and IL-6 mutants. HeLa cells, overexpressing gp130 were stimulated for 45 min with 40nM of each cytokine. Ligand uptake into endosomal structures was co-localized with EEA1. Co-localization of ligands with EEA1 endosomes are highlighted in zoomed area. (e) Quantification of Ligand binding/uptake. Mean fluorescence intensity of DY547-conjugated IL6 variants colocalizing with EEA1 positive structures quantified using the Volocity 3D Image Analysis software (PerkinElmer).

Interestingly, although we detected strong STAT3 activation by C7 and A1 variants (Fig. 2 a-d), their ability to dimerize gp130 in live cells was significantly compromised (Fig. 3 b). Based on this, we speculated that complexes formed by these variants were short-lived and escaped detection by single molecule tracking. It is accepted that cytokine-receptor complex rapidly traffic to intracellular compartments, where they can be degraded or recycled (Gonnord et al., 2012). Recently, it has been proposed that endosomes could act as signaling hubs, helping to sustain low-affinity cytokine-receptor dimers by enhancing the local receptor density (Gandhi et al., 2014). Thus, we asked whether the biased signaling program engaged by the three IL-6 variants resulted from differential receptor trafficking. To test this hypothesis, we fluorescently labelled HyIL-6 and the three IL-6 variants and followed their receptor-mediated internalization by confocal imaging. Importantly, the dye-conjugated IL-6 variants induced dimerization of endogenous gp130 in HeLa cells, confirming their functionality (Sup. Fig 2 c). HeLa cells were incubated for 30 min with the labelled cytokines and their internalization monitored by confocal microscopy. Anti-EEA1 antibodies were used to labelled early endosomes. As shown in Fig. 3 d-e, we detected high levels of labelled Hy-IL6 in intracellular compartments that partially co-localized with EEA1 early endosome marker. Yet, much weaker levels of labelled Mut3 were detected, and no fluorescence was detected for C7 and A1 variants, despite moderate overexpression of gp130. Overall, these data suggest that decreases in gp130 binding affinity impact IL-6-gp130 intracellular traffic, which ultimately could explain the biased signaling programs engaged by the IL-6 variants.

### Gp130 internalization blockages differentially controls STAT1 activation

IL-6 stimulation drives proteasomal degradation of gp130 (Tanaka et al., 2008). Next, we studied whether stimulation of HeLa cells with the three IL-6 variants produced different levels of gp130 degradation. HeLa cells were stimulated with saturating concentrations of HyIL-6 or the different IL-6 variants for 3 hr in the presence of cycloheximide to prevent new protein synthesis. As shown in Fig. 4 a, while HyIL-6 induced a strong decrease in gp130 levels, the ability of the three IL-6 variants to degrade gp130 was significantly reduced. These results show that activation of signaling pathways leading to receptor degradation can be decoupled from STAT1/3 activation by modulating cytokine-receptor complex half-life. Indeed, while Mut3 activates STAT1 and STAT3 to the same extent as HyIL-6, it induced substantially lower degradation of gp130 (Fig. 4 a).

**Figure 4:**
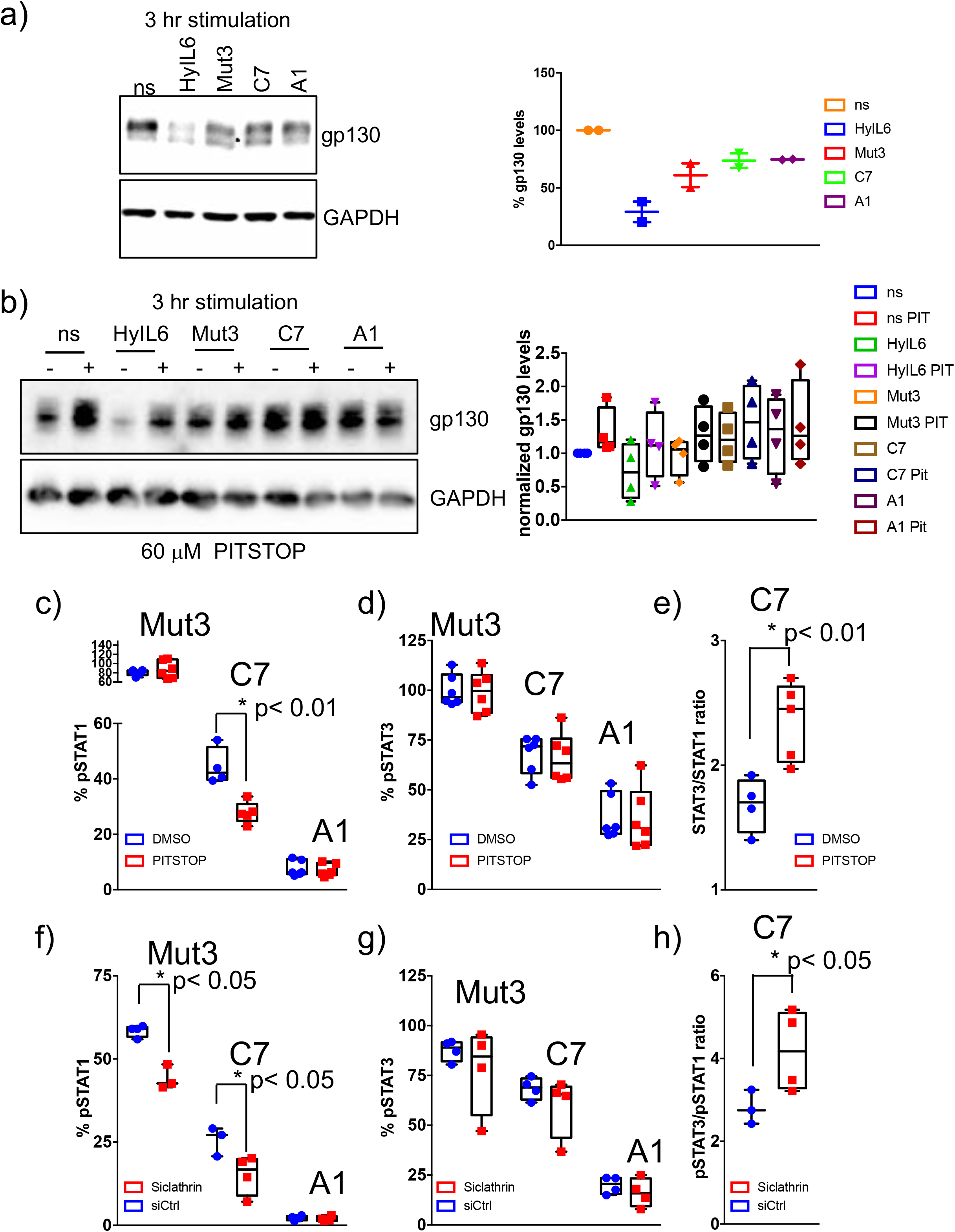
Role of receptor internalization in STAT activation by IL-6 variants. (a) HeLa cells were stimulated with saturating concentrations (100 nM) of the indicated IL-6 ligands for three hours. Levels of gp130 were measured by western blotting using a gp130 specific antibody and quantified via ImageJ software. Data are mean +/− SD of two independent experiments. (b) Hela cells were pre-incubated with 60 μM Pitstop or DMSO for 30 min and then stimulated with saturating concentrations (100 nM) of the indicated IL-6 ligands for 3 hours. Levels of gp130 were measured by western blotting using a gp130 specific antibody and quantified via ImageJ software. Data are mean +/− SD of four independent replicates. (c-e) HeLa cells preincubated for 30 min with Pitstop or DMSO were stimulated with saturating concentrations (100 nM) of the indicated IL-6 ligands for 15 min and levels of STAT1 (c) and STAT3 (d) activation were measured by phospho-Flow cytometry. Data are mean +/− SEM from three independent replicates, each performed in duplicate. The pSTAT3/pSTAT1 ratio calculated from these studies is plotted in (e). (f-h) HeLa cells were transfected with either control siRNA or clathrin specific siRNA. After 48 hours cells were stimulated with saturating concentrations (100 nM) of the indicated IL-6 ligands for 15 min and the levels of STAT1 (f) and STAT3 (g) activation were measured by phospho-Flow cytometry. Data are mean +/− SEM from two independent replicates, each performed in duplicate. The pSTAT3/pSTAT1 ratio calculated from these studies is plotted in (h).

To investigate whether the defective gp130 internalization/degradation induced by the IL-6 variants was at the basis of their biased signaling program, we blocked gp130 internalization by incubating HeLa cells with Pitstop, a well-known clathrin inhibitor. Pitstop-treated HeLa cells exhibited increased basal levels of gp130, possible due to blockage of steady-state gp130 traffic (Fig. 4 b). Moreover, HyIL-6-induced gp130 degradation was blocked confirming the clathrin-dependent internalization of gp130 (Fig. 4 b) (Tanaka et al., 2008). In agreement with Fig. 4 a, the three IL-6 variants did not induce gp130 degradation in the presence/absence of the clathrin inhibitor (Fig. 4 b). We next measured STAT1 and STAT3 phosphorylation levels induced by Mut3, C7 and A1 variants in HeLa cells pre-incubated with Pitstop (Fig. 4 c-e). As shown in Fig. 4 d, the STAT3 phosphorylation levels induced by the three IL-6 variants did not change when gp130 internalization was blocked. STAT1 activation by Mut3 was not affected and A1 failed to activate STAT1 as shown in previous experiments (Fig. 4 c). STAT1 phosphorylation levels induced by the C7 variant on the other hand were significantly downregulated in the presence of Pitstop (Fig. 4 c), which ultimately led to a more pronounced STAT3/STAT1 activation ratio by this variant (Fig. 4 e). We could confirm these data by silencing clathrin in HeLa cells using siRNA (Fig. 4 f-h, Sup. Fig.3 a-b). Clathrin silencing did not affect STAT3 activation (Fig. 4 g), but reduced activation of STAT1 by Mut3 and C7 variants (Fig. 4 f). Overall, these data indicate that translocation of gp130 complexes to intracellular compartments is an important requisite for STAT1, but not STAT3 activation by short-lived IL-6-gp130 complexes.

### STAT1 and STAT3 compete for phospho-Tyrosines in the gp130 intracellular domain

We have shown that trafficking of IL-6/gp130 complexes to intracellular compartments preferentially modulates STAT1 activation. However, why STAT1 activation requires receptor internalization is not well defined. Previous work showed that STAT3, via its SH2 domain, binds with higher affinity than STAT1 to phospho-Tyr on gp130 (Wiederkehr-Adam et al., 2003). We thus postulated that competitive binding of STAT1 and STAT3 for phospho-Tyr on gp130 would result in differential levels of activation of these two transcription factors in the context of short-lived IL-6/gp130 complexes. To test this model, we generated a chimera receptor system, based on the IL-27 receptor complex, to study the influence of the number of phospho-Tyr available in gp130 on ligand-induced STAT1 and STAT3 activation.

IL-27 triggers signaling by dimerizing IL-27Rα and gp130 receptor subunits (Stumhofer et al., 2010). We took advantage of the shared used of gp130 by the two systems and swapped the intracellular domain of IL-27Rα with that of gp130. Additionally, we generated a second receptor chimera (gp130 del-Y), where the intracellular domain of IL-27Rα was swapped with the gp130 intracellular domain containing a deletion after JAK1 binding site, i.e. the box1-2 region. As a result of this, while the first chimera receptor can trigger the potential phosphorylation of eight Tyr, the second chimera can only induce the phosphorylation of four (Fig. 5 a). We then stably transfected all the constructs in RPE1 cells, which do not express IL-27Rα endogenously, but express endogenous levels of gp130 (Fig. 5 b). To ensure that all the RPE1 clones were homogenous and the effects that we see are specific, we compared the responsiveness of the three clones to HyIL-6. As shown in Sup. Fig. 3 c, HyIL-6 induced comparable levels of STAT1 and STAT3 activation in the three clones, strongly arguing that the endogenous gp130, JAK1, STAT1 and STAT3 levels in the three clones were identical. In response to IL-27, the three clones produced very similar STAT3 activation levels, suggesting that STAT3 activation is very efficient and only requires a minimal set of phospho-Tyr available to reach its activation peak (Fig. 5 b and Sup. Fig S3 d). However, STAT1 activation levels dropped by more than fifty percent in the gp130del-Y clone (Fig. 5 b and Sup. Fig S3 d), demonstrating that STAT1 activation by gp130 requires a higher number of phospho-Tyr available. To further reinforce this model, we evaluated whether STAT1 activation levels could be rescued by decreasing STAT3 protein levels. For that, HeLa cells were transfected with either scrambled siRNA or siRNA specific for STAT3 and then stimulated with the different IL-5 variants. STAT3 levels were decreased by more than 80% in transfected cells (Sup. Fig. 3 e). As expected, cell lacking STAT3 expression fail to induce STAT3 phosphorylation (Fig. 5 c). While STAT3 silencing did not change the STAT1 activation levels induced by HyIL-6 and Mut3, STAT1 activation was significantly increased upon C7 stimulation, suggesting a competition between STAT1 and STAT3 for Tyr in gp130 (Fig. 5 c and Sup. Fig. 3 d). Overall, our data strongly support a model where ligand-receptor complex half-life and STATs competition for phospho-Tyr on cytokine receptor intracellular domains maintain a tight equilibrium that allow cells to fine tune their signaling output upon cytokine stimulation.

**Figure 5:**
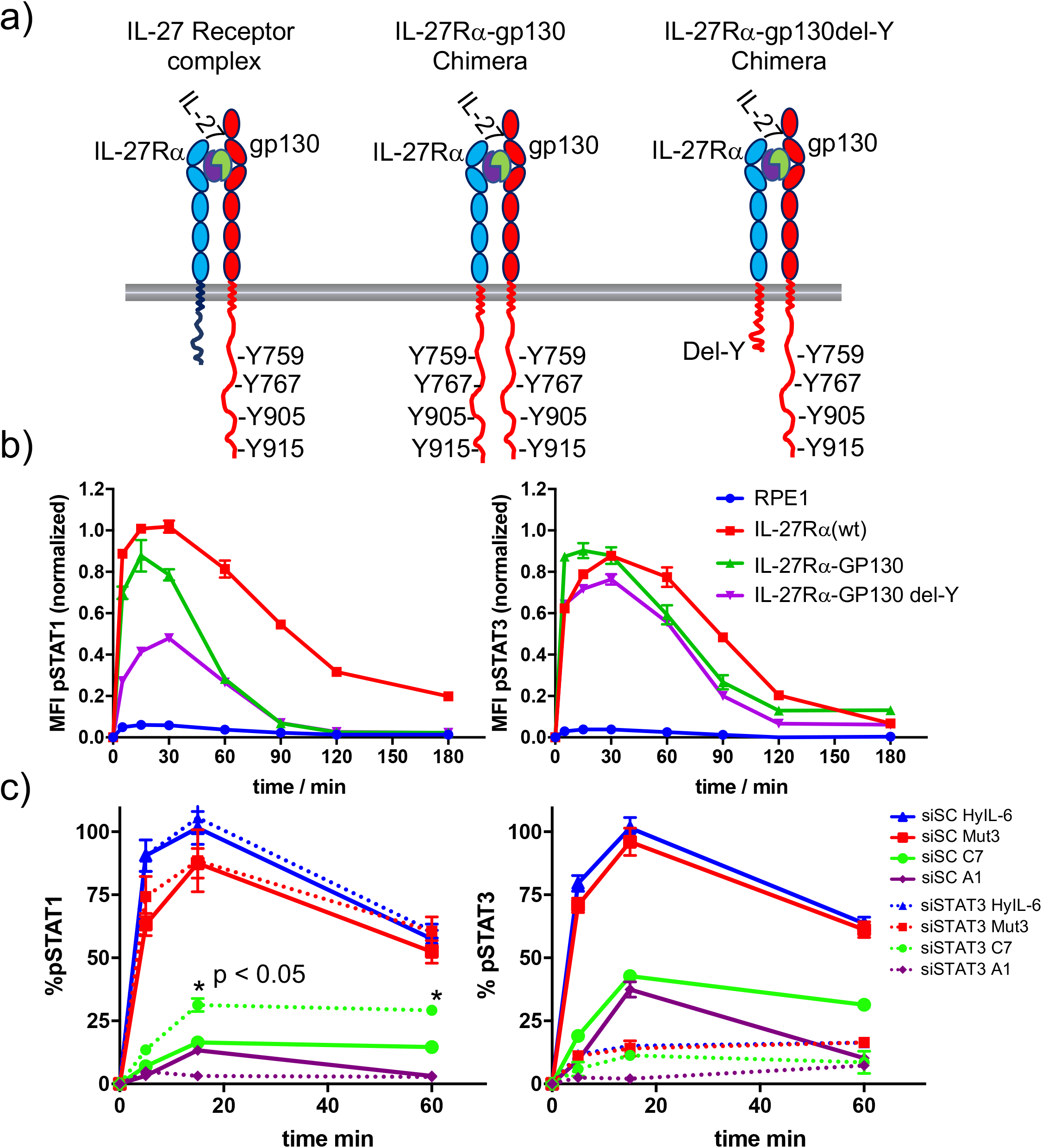
Correlation between number of P-Tyr in gp130 ICD and STATs activation. (a) Schematic representation of the different chimera receptors designed for this study. IL-27Rα intracellular domain was swapped for that of gp130 or a truncated version of the latter lacking all Tyr residues after the box1/2 region. This results in a receptor chimera complex able to engaged 8 P-Tyr and another one able to engage only 4 P-Tyr. RPE1 clones stably expressing the different receptor chimera constructs were generated. (b) Stable RPE1 clones were stimulated with saturating concentrations of IL-27 for the indicated times and the levels of STAT1 (left panel) and STAT3 (right panel) activation were measured by Phospho-Flow cytometry. Data are mean +/− SEM from three independent replicates, each performed in duplicate. (c) Hela cells were transfected with either control siRNA or siRNA targeting STAT3. After 48 hours transfected cells were stimulated with saturating concentrations of the different IL-6 ligands for the indicated times and the levels of STAT1 (left panel) or STAT3 (right panel) activation were measured by Phospho-Flow cytometry. Data are mean +/− SEM from three independent replicates, each performed in duplicate.

### IL-6 variants induce graded STAT3 transcriptional responses

Our data clearly indicate that cytokine-receptor complex half-life instructs biased signaling output by cytokine receptors. However, whether the observed changes in signaling ultimately translate into proportional gene expression changes and bioactivities is not clear. To investigate the immediate effects of “biased” IL-6 signaling input on transcriptome of immune cells, we have generated global transcriptional profiles elicited by the three IL-6 variants in human Th1 cells. First, we performed signaling experiments in human Th1 T cells, to confirm signaling biased by the IL-6 variants in cells expressing gp130 and IL-6Rα receptor subunits simultaneously. Purified human CD4 T cells were activated through its T cell receptor (TCR) *in vitro* and expanded in Th-1 polarizing conditions for five days before they were stimulated with saturating doses of HyIL-6 or the three IL-6 variants. As in HeLa cells, Mut3 activated STAT1 and STAT3 to the same extent as HyIL-6 in Th1 cells (Sup. Fig. 4 a-b). C7 and A1 activated STAT1 and STAT3 to different extents with C7 activating 60% STAT1 and 85% STAT3 when compared to HyIL-6, and A1 activating 50% STAT1 and 70% STAT3 when compared to HyIL-6 (Sup. Fig. 4 a-b). These resulted in an increased STAT3/STAT1 activation ratio by C7 and A1 variants (Sup. Fig. 4 c).

Accordingly, to quantify its effects on gene expression, Th1 cells were stimulated with saturated concentrations of the three IL-6 variants and HyIL-6 for six hours to ensure that the entire cell population respond uniformly to the respective cytokine stimulation and their gene expression program analyzed by RNA-seq studies. We detected the upregulation of 23 genes in response to all four IL-6 variants (Fold change > 1.5, FDR < 0.05, RPKM >4; Fig. 6 a-b), which were all classical STAT3-induced genes, validating the RNA-seq study. Importantly, most target genes showed a graded increase in the rate of transcription as a function of increasing STAT3 activity as exhibited by the three IL-6 variants (pSTAT3 levels; hyIL-6=Mut3>C7>A1). However, the magnitude of transcriptional outputs differ widely from gene to gene, with some genes achieving maximal transcript levels even at low STAT3 levels (Fig. 6b). For instance, while Mut3 gene signature resembles that of HyIL-6, C7 and A1 variants gene signatures exhibited a graded response, with some genes induction decreased by fifty percent (e.g. *SOCS3* and *BCL3*) and other genes expression barely affected when compared to HyIL-6 (e.g. *JAK3, ANK3, PIM2*). We further confirmed these observations by qPCR studies (Sup. Fig.4 d-g). Overall these results suggest that IL-6-induced genes are differently sensitive to corresponding changes in nuclear STAT3 levels, which could provide the cell with the necessary flexibility to fine-tune its responses to wide-range of cytokines levels.

**Figure 6:**
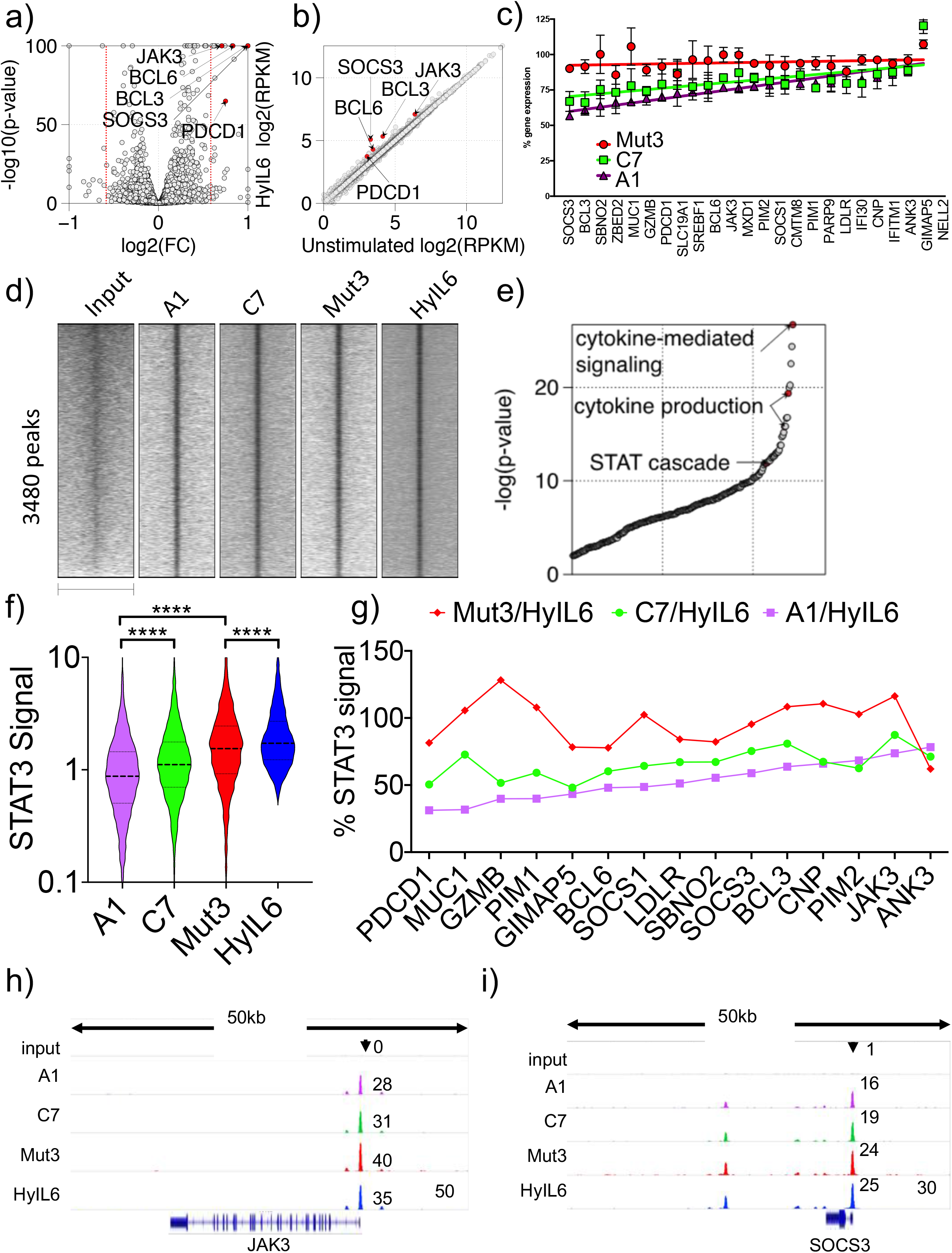
Transcriptional program elicited by the different IL-6 variants. (**a**) volcano plot showing significant genes differently expressed in Th1 cells after 6 hr stimulation with HyIL6. The red dash lines demark fold change = 1.5. (**b**) scatter plot showing mean gene expression values (n=3) before (X-axis) and after HyIL6 stimulation (Y-axis). Top five differently expressed genes are highlighted. (**c**) plot showing the normalized gene expression relative to HyIL6 stimulation for each indicated stimulation. 23 differently expressed genes after HyIL6 stimulation are shown. The regression lines are highlighted. The data in **a-c** are from three independent donors. (**d**) heatmap showing signal intensity of STAT3 bound regions (5kb centred at peak summit) for indicated stimulations. Peaks are identified by comparing Hy-IL6 stimulation and input. (**e**) shown are GO biological pathways ranked by p-value that are enriched in genes with adjacent STAT3 binding. (**f**) violin plot showing the signal intensity of all peaks (200bp regions centred at peak summit) after each stimulation. P values are determined by two-tailed Wilcoxon tes (**** P<0.0001). (**g**) shown are relative signal intensity of STAT3 peaks near select genes. Select are 15 differently expressed genes with adjacent STAT3 binding sites. (**h-i**) STAT3 binding at JAK3 (**h**) and SOCS3 (**i**) gene loci.

Next, to investigate how IL-6-induced STAT3 sites within the genome orchestrate the observed graded gene expression response, we measured global STAT3 binding profiles by ChIP-seq and compared the transcriptional activity of its target genes. Specifically, given that IL-6 variants induced different levels of STAT3 phosphorylation, we quantified genome-wide STAT3 binding sites in Th-1 cells as a function of gradient STAT3 activation by the IL-6 variants. As expected, IL-6 stimulation led to STAT3 binding to 3480 genomic loci (Fig. 6 d), which were localized near classical STAT-associated genes (Fig. 6 e). We could detect significant changes in STAT3 binding intensity in response to the different IL-6 variants, which correlated with their STAT3 activation levels (Fig. 6 f). Of note, although ChIP-seq data identified many genome-wide IL-6-induced STAT3 binding sites, only a handful of those STAT3-target genes (23 transcripts) were upregulated in Th1 cells, suggesting additional mechanisms by which IL-6-induced STAT3 influences gene expression programs. Moreover, when we examined STAT3 bound regions near genes upregulated by IL-6 stimulation (Fig. 6 c), we observed a similar trend to that observed in the RNA-seq studies, i.e. STAT3 binding intensities were more different in those genes differentially regulated by the IL-6 variants (eg. *BCL3* and *SOCS3*), and more similar in genes equally regulated by all four ligands (e.g. *JAK3* and *PIM2*) (Fig. 6 g-i and Sup. Fig. 4 h-i). Interestingly, *SOCS3* and *BCL3* that were among the most differentially expressed IL-6-induced genes, contain multiple STAT3 binding sites (Supplemental Table 1), which may enable IL-6 to produce graded transcriptional outputs among its target genes. By contrast, STAT3 target genes with 1 or 2 binding sites at the gene promoter become saturated at relatively low levels of STAT3 transcriptional activation. This suggests that genes with multiple STAT3 binding sites would be more sensitive to changes in STAT3 signaling levels compared to gene with a single STAT3 binding site. Collectively, our data indicates that IL-6 variants result in graded STAT3 binding and transcriptional responses.

### IL-6 variants induce immuno-modulatory activities with different efficiencies

IL-6 is a highly immune-modulatory cytokine, contributing to the inflammatory response by inducing differentiation of Th-17 cells and inhibition of Treg and Th-1 cells (Heink et al., 2017; Jones et al., 2010; Kimura and Kishimoto, 2010; Louten et al., 2009) (Fig. 7 a-c). We next asked whether these three activities would be uniformly affected by the biased signaling programs engaged by the three IL-6 variants. For that, we cultured resting human CD4 T cells in Th-17, Th-1 and Treg polarizing conditions in the presence/absence of the different IL-6 variants. As shown in Fig. 7, the three variants induced responses that parallel their STAT activation potencies (Fig. 7 d-f). However, not all three activities were equally engaged by the three IL-6 variants. While all variants induced differentiation of Th-17 cells to some extent (Fig 7 d and Sup. Fig. 5), C7 and A1 variants struggle to inhibit differentiation of Treg and Th-1 cells, with C7 eliciting some inhibition and A1 failing in both cases (Fig. 7 e-f and Sup. Fig. 5). This is better represented in Fig. 7 g, where a triangular illustration is used to show that Mut3 is equally potent in inducing the three activities, producing an equilateral triangular shape. C7 and A1 on the other hand produced non-equilateral triangular shapes, exhibiting different induction efficiencies of the three bioactivities. Overall, these results show that not all cytokine bioactivities require the same signaling threshold, and that by modulating cytokine-receptor binding parameters and cytokine-induced signaling programs, we can decouple or at least biased these responses.

**Figure 7:**
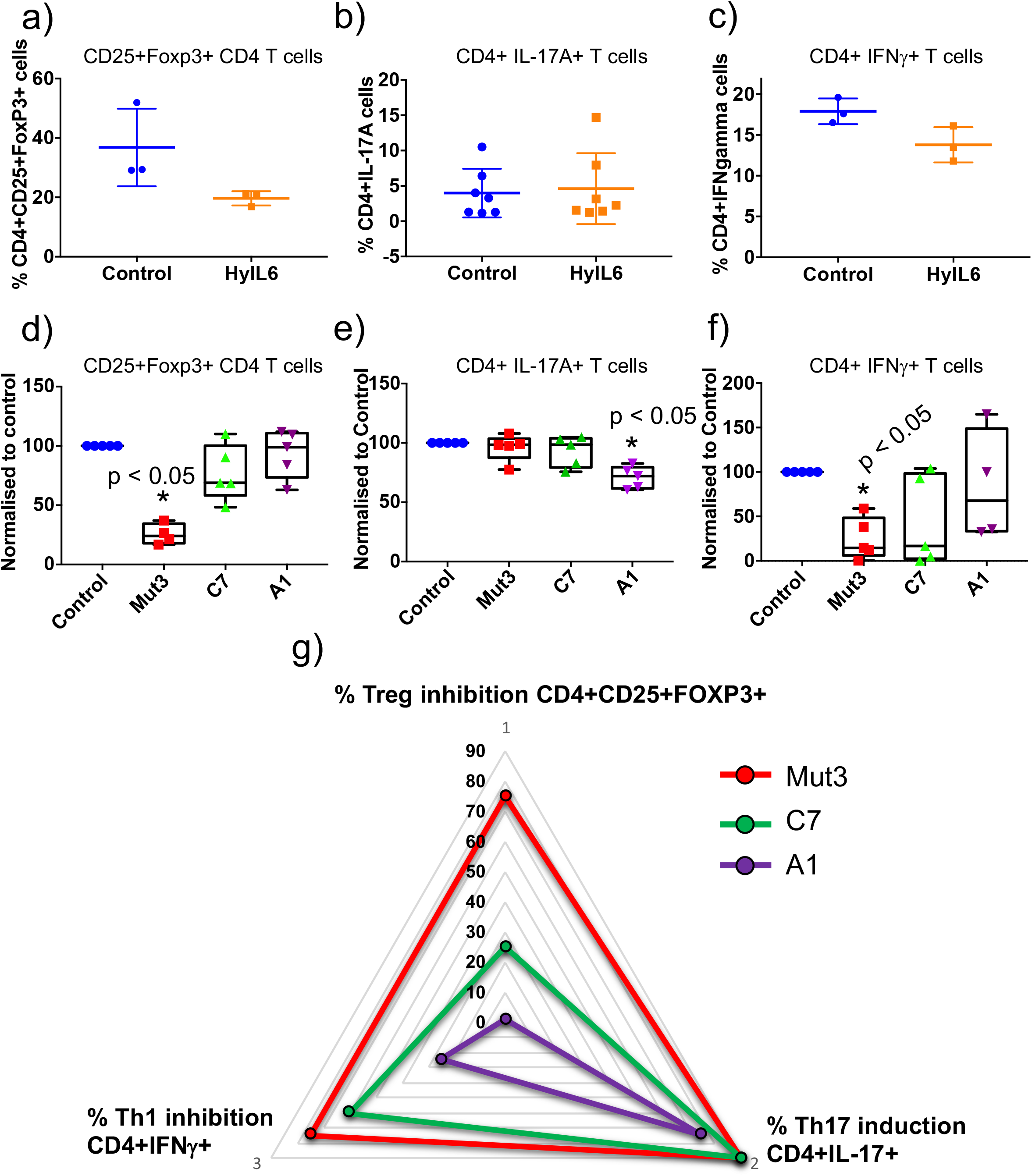
Immuno-modulatory activities trigger by the different IL-6 variants. (a) Human CD4 T cells were isolated from whole PBMCs and treated with Treg polarizing conditions in the presence of saturating concentrations of the different IL-6 variants for five days. Percentage of Treg cells were calculated by counting number of events in the CD4+CD25+FoxP3+ population obtained by flow cytometry. The control condition was defined as 100 % response and the other conditions normalized accordingly. Data are mean +/− SEM from five independent replicates. (b) Human CD4 T cells were isolated from whole PBMCs and treated with Th-17 polarizing conditions in the presence of saturating concentrations of the different IL-6 variants for fourteen days. Percentage of Th-17 cells were calculated by counting number of events in the CD4+IL-17A+ population obtained by flow cytometry. The control condition was defined as 100 % response and the other conditions normalized accordingly. Data are mean +/− SEM from five independent replicates. (c) Human CD4 T cells were isolated from whole PBMCs and treated with Th-1 polarizing conditions in the presence of saturating concentrations of the different IL-6 variants for five days. Percentage of Th-1 cells were calculated by counting number of events in the CD4+IFNγ+ population obtained by flow cytometry. The control condition was defined as 100 % response and the other conditions normalized accordingly. Data are mean +/− SEM from four independent replicates. (d) Triangular representation of data from (a-c). As the affinity for gp130 decreases (C7 and A1 variants) the different IL-6 activities are differentially affected with Th-17 differentiation being the most robust activity to changes in affinity and Treg inhibition being the most sensitive activity.

## DISCUSSION

In this study we have engineered IL-6/gp130 binding kinetics to modulate signaling output and decouple IL-6 functional pleiotropy. Two main findings arise from our study: (1) Intracellular traffic dynamics of cytokine-receptor complexes and STATs binding affinities for phospho-Tyr on cytokine receptor intracellular domains (ICD) act synergistically to define signaling potency and identity, and (2) cells exhibit different gene induction thresholds in response to cytokine partial agonism, which allow them to modulate their responses. The current work, together with previous studies describing signaling tuning in other cytokines systems (Ho et al., 2017; Kim et al., 2017; Moraga et al., 2015b,Mitra et al., 2015), outline a general strategy to design cytokine partial agonists and decouple cytokine functional pleiotropy by modulating cytokine-receptor binding kinetics.

All our IL-6 variants must dimerize gp130 to some extent, since they all trigger signaling. However, our single molecule TIRF data show that low affinity variants C7 and A1 struggle to promote detectable gp130 dimerization. These data suggest that non-detectable short-lived IL-6/gp130 complexes can partially engage signaling, but fail to trigger a full response, evoking a kinetic-proof reading model. A kinetic-proof reading model has been previously proposed for other ligand-receptor systems, including the T cell receptor system (TCR) (McKeithan, 1995) and more recently Receptor Tyrosine Kinases (RTKs) (Zinkle and Mohammadi, 2018). In these two systems, changes on ligand-receptor complex half-life induce phosphorylation of different Tyr pools in the receptors ICDs, ultimately recruiting and activating different signaling effectors (Acuto et al., 2008; Lemmon and Schlessinger, 2010). More recently, this model was used to explain biased signaling triggered by an EPO mutant (Kim et al., 2017). However, cytokine receptors differ significantly from RTKs and the TCR. While in these latter receptor systems, activation of different signaling effectors is clearly assigned to phosphorylation of specific Tyr in their ICDs, this is not generally true for cytokine receptors. Often only one or two Tyr in the cytokine receptor ICDs are required for signal activation (Cheng et al., 2011; Schmitz et al., 2000; Zhao et al., 2008). In this context, it is difficult to reconcile the canonical kinetic-proof reading model with signaling biased triggered by cytokines.

How can short-lived cytokine-receptor complexes engage different signaling effectors that compete for a single phospho-Tyr? Our study now provides new molecular evidences that shed light into this question. We showed, using chimera receptors and siRNA approaches, that binding affinity of STAT proteins for phospho-Tyr in receptors ICDs defines signaling amplitude and identity by cytokines. Importantly, this is not the first evidence of STATs competing for phospho-Tyr. IFNα2 activates all STATs molecules, which can be abrogated by a single Tyr mutation in the IFNAR2 ICD, suggesting STAT competition (Zhao et al., 2008). Moreover, modulation of STAT protein levels has been described to change signaling specificity by cytokines. IFNγ priming, which result in enhanced STAT1 protein levels, shift the IL-10 response from STAT3 activation to STAT1 activation (Herrero et al., 2003). In cells lacking STAT3, IL-6 switches to STAT1 activation, producing IFNγ-like responses (Costa-Pereira et al., 2002). These observations strongly argue in favour of a model where STATs compete for Tyr in receptors ICDs, thus making them sensitive to changes in complex halflife. STATs binding with low affinity to phospho-Tyr would require more stable cytokine-receptor complexes and higher ligand doses to reach maximal activation. In agreement with this model, a previous study reported STAT1 and STAT3 binding with different affinities to phospho-Tyr in gp130 ICD (Wiederkehr-Adam et al., 2003).

Initially thought to contribute to cytokine signaling shutdown, the endosomal compartment has emerged in recent years as a signaling hub, not only in cytokines (Becker et al.; Bulut et al.; Claudinon et al., 2007; German et al.; Keeler et al., 2007; Shah et al., 2006) but also in other ligand-receptor systems (Villasenor et al., 2016). Previous studies showed that activated JAK and STATs molecules are found in endosomes upon cytokine stimulation (Payelle-Brogard and Pellegrini, 2010), suggesting that cytokine-receptor complexes exhibit a signaling continuum from the cell surface to intracellular compartments. In agreement with this model, recent studies showed that cytokine-receptor complexes traffic to the endosomal compartment, where they are stabilized, contributing to signaling fitness (Gandhi et al., 2014; Moraga et al., 2015a). Only short-lived complexes that fail to traffic to intracellular compartments trigger diminish signaling output. Our data fully support this model and expand it by showing that short-lived complexes that fail to traffic to the endosomal compartment engage a biased signaling program. Low affinity IL-6 variants, which did not induce internalization of gp130, activated more efficiently STAT3 than STAT1. The stabilization of short-live complexes in endosomes provide the extra time necessary to activate secondary pathways that engage the receptor with lower affinity. An open question pertains to whether the intracellular localization of STATs influence signaling by long- or short-live cytokine receptor complexes. Early studies described the localization of STAT3 in intracellular membranes, while other STATs association with intracellular membranes is not so well described (Shah et al., 2006). Whether signaling by cytokines can be further engineered by modulation of the intracellular localization of STATs requires further investigation.

In the current study, we show that different genes downstream of IL-6 signaling exhibit different thresholds of activation that can be exploited by IL-6 partial agonists to decouple IL-6 immunomodulatory activities. How these activation thresholds are established is not clear. STAT proteins face two points where the law of mass action influences their responses the most. The first one pertains to the binding of STATs to phopho-Tyr in the receptors ICDs, and as discussed above, contributes to define signaling potency and identity by cytokine-receptor complexes. The second point is found when activated STATs bind specific GAS sequence motifs in the promoters of responsive genes. GAS sequences, although conserved, exhibit degrees of degeneracy that allow them to bind STATs molecules with different affinities (Bonham et al., 2013; Ehret et al., 2001; Horvath et al., 1995). In addition, different number of GAS sequences are found in different responsive promoters. In principle, the combination of STAT binding affinities for GAS sequences and the number of GAS sequence present in the promoters could generate different gene induction thresholds. In agreement with this model, we identified chromatin regions through our STAT3 Chip-Seq studies, that bound STAT3 with different efficiencies, with regions where STAT3 binding was diminished by changes in STAT3 activation levels, and regions where efficient STAT3 binding was detected in all conditions tested. When we analysed the number of GAS sequences and their motifs under those regions, we could detect that genes that were more sensitive to changes in STAT3 phosphorylation presented higher number of GAS motifs than those more resistant. Overall our data support a kinetic-proof reading model for cytokine signaling, whereby cytokine-receptor dwell time and STAT binding affinities for phospho-Tyr on receptors ICDs define potency and identity of cytokine signaling signatures. As the number of STAT molecules activated by partial agonists increases, additional GAS binding motifs are engaged in promoters with multiple GAS binding sites, triggering the induction of graded gene expression responses. In principle, engineering of cytokine-receptor binding kinetics could rescue cytokine-based therapies, by decreasing cytokine functional pleiotropy and toxicity.

## ACKNOWLEDGMENTS

We thank members of the Moraga, Mitra and Kazemian laboratories for helpful advice and discussion. We thank Dynamic Biosensors for their help characterizing IL-6 variants binding properties. This work was supported by the StG, LS6, ERC-206-STG grant (I.M.) and by SFB 944 (P8, J.P.)

## AUTHOR CONTRIBUTIONS

M.F.J., W.S. and M.I. conceived the project; M.F.J., W.S., W.L., K.M., M.S., M.I. wrote the manuscript; M.I. performed the engineering studies. M.F.J., W.S., P.E., L.J. and C.G. performed signaling and cellular experiments; W.S. H.M. and P.J. designed and performed single-particle microscopy experiments; M.F.J. W.L., K.M. and M.S. designed, performed and analysed Chip-Seq and RNA-Seq studies.

## MATHERIAL AND METHODS

### Protein expression and purification

Human IL-6 wild type and IL-6 variants were cloned into the pAcGP67-A vector (BD Biosciences) in frame with an N-terminal gp67 signal sequence and a C-terminal hexahistidine tag, and produced using the baculovirus expression system, as described in (LaPorte et al., 2008). Baculovirus stocks were prepared by transfection and amplification in *Spodoptera frugiperda* (*Sf9*) cells grown in SF900II media (Invitrogen) and protein expression was carried out in suspension *Trichoplusiani ni* (High Five) cells grown in InsectXpress media (Lonza). Following expression, proteins were captured from High Five supernatants after 48 hrs by nickel-NTA agarose (Qiagen) affinity chromatography, concentrated, and purified by size exclusion chromatography on a Enrich SEC 650 1×300 column (Biorad), equilibrated in 10 mM HEPES (pH 7.2) containing 150 mM NaCl. Recombinant cytokines were purified to greater than 98% homogeneity. For biotinylated gp130 expression, gp130 ectodomain (SD1-SD3, amino acids 23-321) was cloned into the pAcGP67-A vector with a C-terminal biotin acceptor peptide (BAP)-LNDIFEAQKIEWHW followed by a hexahistidine tag. Purified Gp130 was in vitro biotinylayed with BirA ligase in the presence of excess biotin (100 μM). HyIL-6 was site-specifically labeled via an ybbR-tag by enzymatic phosphopantetheinyl-transfer using coenzyme A conjugates as described previously (Waichman et al., 2010). For site-specific fluorescence labeling of IL-6 variants with different fluorochromes, an accessible cysteine was introduced at the C-terminal of the cytokine and cloned in the pAcGP67-A vector as described above. Labeling of purified proteins was carried out with excess DY 647 and DY 547 maleimide, repectively in the presence of 50 μM TCEP.

### Plasmid constructs

For single molecule fluorescence microscopy, monomeric non-fluorescent (Y67F) variant of eGFP (“mXFP”) was N-terminally fused to gp130. This construct was inserted into a modified version of pSems-26 m (Covalys) using a signal peptide of Igk. The ORF was linked to a neomycin resistance cassette via an IRES site. A mXFP-IL-27Rα construct was designed likewise. The chimeric construct mXFP-IL-27-Rα (ECD)-gp130(ICD) was a fusion construct of IL-27Rα (aa 33-540) and gp130 (aa 645-918). For mXFP-IL-27-Rα(ECD)-gp130(ICD)-Box1/2 Δ the ICD of gp130 was truncated downstream of the JAK1 binding motif (aa 645-705).

### Cell lines and media

HeLa cells were grown in DMEM containing 10% v/v FBS, penicillin-streptomycin, and L-glutamine (2 mM). RPE1 cells were grown in DMEM/F12 containing 10% v/v FBS, penicillin-streptomycin, and L-glutamine (2 mM). HepG2 cells and Ba/F3-gp130 (Gearing et al., 1994) cells were cultured in DMEM containing 10% v/v FBS, and penicillin-streptomycin. Viability of Ba/F3-gp130 cells was determined as described previously (Garbers et al., 2011). Human T-cells were cultivated in RPMI supplemented with 10% v/v FBS, penicillin-streptomycin and cytokines for proliferation/differentiation (see below). RPE1 cells were stably transfected by mXFP-IL-27-Rα and the chimeric constructs by PEI method according to standard protocols. Using G418 selection (0.6 mg/ml) individual clones were selected, proliferated and characterized. For comparing receptor cell surface expression levels, cells were detached using PBS+5mM EDTA, spun down (300g, 5 min) and incubated with αGFP-nanobody^Dy647^ (10nM, 15 min on ice). After incubation, cells were washed with PBS and run on cytometer.

### CD4+ T cell purification

Peripheral blood mononuclear cells (PBMCs) of healthy donors were isolated from buffy coat samples (Scottish Blood Transfusion Service) by density gradient centrifugation according to manufacturer’s protocols (Lymphoprep, STEMCELL Technologies). From each donor, 100×10^6^ PBMCs were used for isolation of CD4+ T-cells. Cells were decorated with anti-CD4+^FITC^ antibodies (Biolegend, #357406) and isolated by magnetic separation according to manufacturer’s protocols (MACS Miltenyi) to a purity >98% CD4+.

### Flow cytometry staining and antibodies

For measuring dose-response curves of STAT1/3 phosphorylation (either TH1 cells or HeLa/RPE1 clones), 96-well plated were prepared with 50μl of cell suspensions at 2×10^6^ cells/ml/well for TH1 and 2×10^5^ cells/ml/well for HeLa/RPE1. RPE1 cells were detached using Accutase (Sigma). Cells were stimulated with a set of different concentrations to obtain dose-response curves. To this end cells were stimulated for 15 min at 37°C with the respective cytokines (mIL27sc or hypIL6) followed by PFA fixation (2%) for 15 min at RT.

For kinetic experiments, cell suspensions were stimulated with a defined, saturating concentration of cytokines (2nM mIL27sc, 10nM hypIL6, 100nM IL-6 mutants) in a reverse order so that all cell suspensions were PFA-fixed (2%) at the same time.

#### Permeabilization, fluorescence barcoding and antibody staining

After fixation (15 min at RT), cells were spun down at 300g for 6 min at 4°C. Cell pellets were resuspended and permeabilized in ice-cold methanol and kept for 30 min on ice. After permeabilization cells were fluorescently barcoded according to (Krutzik and Nolan, 2006). In brief: using two NHS-dyes (PacificBlue, #10163, DyLight800, #46421, Thermo Scientific), individual wells were stained with a combination of different concentrations of these amino-reactive dyes. After barcoding, cells can be pooled and stained with anti-pSTAT1^Alexa647^ (Cell Signaling Technologies, #8009) and anti-pSTAT3^Alexa488^ (Biolegend, #651006) at a 1:100 dilution in PBS+0.5%BSA. T-cells were also stained with anti-CD8^AlexaFlour700^ (Biolegend, #300920), anti-CD4^PE^ (Biolegend, #357404), anti-CD3^BrilliantViolet510^ (Biolegend, #300448). Cells were probed at the flow cytometer (Beckman Coulter, Cytoflex S). Individual cell populations were identified by their barcoding pattern and mean fluorescence intensity (MFI) of pSTAT1^647^ and pSTAT3^488^ was measured for all individual cell populations.

### Western blotting protocol

Cells were rinsed in ice-cold PBS then lyzed in NP40 lysis buffer (1% NP40, 50 mM Tris-HCl pH 8.0, 150 mM NaCl) plus protease inhibitor cocktail (Pierce), 5mM sodium fluoride, 2mM sodium orthovanadate and 0.2mM PMSF incubating on ice for 15 min. Lysates were cleared by centrifugation at 20,000 g for 15 min at 4°C then protein concentrations determined using Coomassie Protein Assay Kit (Thermo Scientific, UK). For each sample, 30 μg of total protein were separated on 7% Bis-Tris polyacrylamide gels in SDS running buffer then blotted onto Protran 0.2mM Nitrocellulose (GE Healthcare, UK). Membranes were probed with 1:1000 dilution of the appropriate primary antibody (mouse anti-gp130; Santa Cruz sc376280), rabbit anti-Clathrin (Biolegend, 813901), STAT3 (Cell Signaling Technologies, #9139), P-STAT3 (Y705, Biolegend, #651006) P-STAT1 (Y701, Cell Signaling Technologies, #8009) or 1:5000 dilution mouse anti-GAPDH (Cell Signaling Technologies, #2118), then 1:5000 dilution of donkey anti-rabbit-HRP (Stratech, 711-035-152-JIR) or donkey anti-mouse-HRP (Stratech, 715-035-150-JIR) as the secondary antibody. Immobilon Western Chemiluminescent HRP substrate (Millipore, UK) was used for visualization.

### siRNA Silencing

HeLa cells were seeded at 2×10^5^ cells per well in a 6 well plate and transfected with Clathrin siRNA (Oligo 1:AGGUGGCUUCUAAAUAUCAUGAACA; Oligo 2: GAAUGUUUACUGAAUUAGCUAUUCT sequences; from IDT Technology), STA3 siRNA (CAACAUGUCAUUUGCUGAA) or non-targeting siRNA (UGGUUUACAUGUCGACUAA) as a control (Dharmacon) using DharmaFect 1 transfection reagent (Dharmacon, Cat#T2001-02) following the manufacturer instructions. 48 hours later cells were treated as indicated and samples were prepared for immunoblotting analysis to check the level of gene knock-down and for FACS (STAT3).

For FACS analysis, cells were stimulated with HyIL6 20nM for 15 minutes, fixed with 2% formaldehyde (Thermo), permeabilised with methanol 100% for 20 minutes at +4°C, and stained for P-Tyr705-STAT3-AF488 (BioLegend, Cat#651006) and P-Tyr701-STAT1-AF647 (CellSignaling, Cat#8009S).

### Assembly, transformation, and selection of the IL-6 library

Yeast surface display protocol was adapted from previously described ones (Boder and Wittrup, 1997). Human IL-6 cDNA was cloned into the yeast display vector pCT302. *S. cerevisiae* strain EBY100 was transformed with the pCT302_IL-6 vector. Generally, yeast were grown in SDCAA media pH: 4.5 for one day, and induced in SGCAA media pH: 4.5 for two days, before undergoing a round of selection. Different concentrations of biotinylated gp130 ectodomains were used to carried out the selections. In initial rounds where gp130-Streptavidin (SA) tetramers were used to select low affinity gp130 binders, tetramers were formed by incubating gp130 and SA coupled to Alexa-647 dye at a ratio of 4:1 gp130:SA for 15 min on ice.

The assembly of the library DNA was carried out using 14 overlapping primers, two of which contained the NDT codon (G,V,L,I,C,S,R,H,D,N,F,Y) used for mutation. The following amino acids were chosen to randomize: D9, E22, R23, K26, Q27, Y30, D33, G34, A37, E109, R112, M116, V120, F124. The PCR product was further amplified, to obtain 50 μg using the primers: 5’-TAGCGGTGGGGGCGGTTCTCTGGAAGTTCTGTTCCAGGGTCCGAGCGGCGGATCCGTACCCCCAGGAGAAGATTCC-3’5’-CGAGCAAGTCTTCTTCGGAGATAAGCTTTTGTTCGCCACCAGAAGCGGCCGCCATTTGCCGAAGAGCCCTCAG-3’

These primers also contained the necessary homology to the pCT302 vector sequence requisite for homologous recombination. Insert DNA was combined with linearized vector backbone pCT302 and electrocompetent *S. cerevisiae* EBY100 were electroporated and rescued, as previously described, forming a library of 3×10^8^ transformants. Selections were performed on this library using magnetic activated cell sorting (MACS, Miltenyi Biotech). The first round of selection was performed with 2×10^9^ cells from the yeast library, approximately 10-fold coverage relative to the number of transformants. Subsequent rounds of selection used 1×10^7^ yeast cells (greater than 10-fold coverage in each round). Fluorescence analysis was performed on a CytoFlex cytometer.

### Determination of binding kinetics by switchSENSE

All measurements were performed on a dual-color DRX^2^ instrument using a standard switchSENSE^®^ chip (MPC2-48-2-G1R1, Dynamic Biosensors GmbH), which provides two differently labeled DNA sequences on each electrode (green fluorescent NL-A48, red fluorescent NL-B48). The chip was functionalized by initial hybridization of streptavidin-cNL-B48 conjugate and bare cNL-A48 DNA (each 200 nM, HE40 buffer, Dynamic Biosensors GmbH). In this way, the red fluorescence yields the signal for the interaction measurement with the target molecule, while the green fluorescence provides an on-spot reference for unspecific effects. In a second step, biotinylated gp130 was injected and captured onto the surface by immobilized streptavidin. To analyze the gp130 – IL-6 interactions, a series of protein concentrations (62 nM–12 μM) was tested. All experiments were performed in HEPES-based running buffer (10 mM HEPES, 140 mM NaCl, 0.05% Tween20, 50 μM EDTA, 50 μM EGTA, pH = 7.4) at 25 °C. For measuring the association, IL-6 variants were injected with a flowrate of 500 μl/min between 60 and 120 s and the absolute fluorescence in static mode was recorded (fluorescence proximity sensing). Dissociations was monitored at the same flow rates (500 μl/min) and varied between 7 min and 3 h depending on dissociation rate constants determined during assay development. After each cycle (analyte concentration), the surface was regenerated and freshly functionalized. Association and dissociation rates were determined by fitting a global mono-exponential model to the raw data.

### qPCR studies

Resting CD4^+^ T cells were labelled with anti-CD4-FiTC antibody (BioLegend, Cat#357406) and isolated from human PBMCs by magnetic activated cell sorting (MACS, Miltenyi) using anti-FiTC microbeads (Miltenyi, Cat#130-048-701) following manufacturer instructions. Subsequently, resting CD4^+^ T cells were activated under Th1 polarizing conditions. Briefly, 10^6^ resting human CD4^+^ T cells per ml were primed for three days with ImmunoCult™ Human CD3/CD28 T Cell Activator (StemCell) following manufacturer instructions in the presence of IL2 (20 ng/ml, Novartis Cat#709421), IL12 (20 ng/ml, BioLegend, Cat#573002) and anti-IL4 (10 ng/ml, BD Biosciences, Cat#554481). Then, cells were expanded in the presence of IL2 (20 ng/ml) and anti-IL4 (10 ng/ml) for another 5 days. Cells were starved without IL2 for at least 24 hours before the stimulation with the different forms of IL6 for 6 hours. Total RNA was isolated using the RNeasy Mini Kit (Qiagen, Cat#74104) and equal amounts of cDNA were synthesised using the iScript cDNA Synthesis Kit (BioRad, Cat# 1708890). 100ng of cDNA were used to assay the expression level of the different genes of interest by qPCR using TB Green Premix Ex Taq II (Takara, Cat# RR820L) in a CFX96 Touch™ Real-Time PCR Detection System (BioRad). GAPDH was amplified as an internal control. The relative quantitation of each mRNA was performed using the comparative Ct method and normalised to the internal control.

Primers for qPCR analysis were:

**GAPDH**

Fw: 5’-ACCCACTCCTCCACCTTTGA-3’ Rv: 5’-CTGTTGCTGTAGCCAAATTGGT-3’,

**SOCS3**

Fw: 5’-GTCCCCCCAGAAGAGCCTATTA-3’ Rv: 5’-TTGACGGTCTTCCGAGAGAGAT-3’,

**BCL3**

Fw: 5’-GAAAACAACAGCCTTAGCATGGT-3’ Rv: 5’-CTGCGGAGTACATTTGCG-3’,

**PIM2**

Fw: 5’-GGCAGCCAGCATATGGG-3’ Rv: 5’-TAATCCGCCGGTGCCTGG-3’

**JAK3**

Fw: 5’-GCCTGGAGTGGCATGAGAA-3’ Rv: 5’-CCCCGGTAAATCTTGGTGAA-3’.

### Chromatin Immunoprecipitation by Sequencing (ChIP-Seq)

In vitro polarized human Th1 cells were expanded in the presence of IL-2 for 10 days and cells were then washed with complete media and rested for 24 hours starvation in the absence of IL-2, these cells were then either not-stimulated (control) or stimulated with IL-6 or different IL-6 variants for 1 hour, cells were then immediately fixed with 1% methanol-free formaldehyde (Formaldehyde 16%, Methanol-Free, Fisher Scientific, PA, USA) at room temperature for 10mn with gentle rocking cells were then washed twice with cold PBS. For each STAT3 ChlP-seq library sample, approximately 10×106 cells were used and the fixed cell palettes were kept at −80°C prior to further processing. The ChIPseq experiments were performed as previously described (PMID:18820682) with some modification as described below. In brief, the frozen cell pellets were thawed on ice and washed once with 1 mL cold PBS by centrifugation at 5000 RPM for 5min, the resulting cell pellets were re-suspended in 500uL of lysis buffer (1X PBS, 0,5% Triton X-100, cOmplete EDTA-free protease inhibitor cocktail, Roche Diagnostics, Basel, Switzerland) and incubated for 10min on ice, followed by a 5min centrifugation at 5000 RPM. Then the pellets were washed once with 1 mL of sonication buffer (1X TE, 1: 100 protease inhibitor cocktail), re-suspended in 750uL of sonication buffer (1X TE,1: 100 protease inhibitor cocktail and 0,5mM PMSF) and sonicated for 20 cycles (on-20sec and off-45sec) on ice using VCX-750 Vibra Cell Ultra Sonic Processor (Sonics, USA). The sonicated lysates were centrifuged 20min at 14000 RPM and the clear lysate supernatants were collected and incubated with 30uL of Protein-A Dynabeads (ThermoFisher, USA) that were pre-incubated with incubated with 10ug of anti-STAT3 antibody (anti-Stat3, 12640S, Cell Signaling Technology) at 4°C overnight with gentle rotation. Next day, the beads were washed 2 times with RIPA-140 buffer (0.1% SDS, 1% Triton X-100, 1mM EDTA, 10mM Tris pH 8.0, 300mM NaCl, 0.1% NaDOC), 2 times with RIPA-300 buffer (0.1% SDS, 1% Triton X-100, 1mM EDTA, 10mM Tris, 300mM NaCl, 0.1% NaDOC), 2 times with LiCl buffer (0.25mM LiCl, 0.5% NP-40, 1mM EDTA, 10mM Tris pH 8.0, 0.5% NaDOC), once with TE-0,2% Triton X-100 and once with TE buffer. Crosslinks were reversed by incubating the bound complexes in 60uL TE containing 4.5uL of 10% SDS and 7.5uL of 20mg/mL of proteinase K (Thermofisher, USA) at 65°C overnight for input samples, we used 6uL of 10% SDS and 10uL of 20mg/mL of proteinase K. Then, the supernatants were collected using a magnet and beads were further washed one in TE 0.5M NaCl buffer. Both supernatants were combined, and DNA was extracted with phenol/chloroform, followed by precipitation with ethanol and re-suspended in TE buffer. The library was constructed following the manufacturer protocol of the KAPA LTP Library Preparation Kit (KAPA Biosystems, Roche, Switzerland). ChIP DNA libraries were ligated with the Bioo scientific barcoded adaptors (BIOO Scientific, Perkin Elmer, USA) with T4 DNA ligase according to KAPA LTP library preparation protocol and the ligated ChIP DNA libraries were purified with 1.8x vol. Agencourt AMPure XP beads and PCR amplified using KAPA hot start High-Fidelity 2X PCR Master Mix and NextFlex index primers (Bioo Scientific, PerkinElmer) for 12 cycle by following thermocycler cycles: 30s hot start at at 98°C, followed by 12 cycle amplification [98°C for 10 sec, 60°C for 30 sec and 72°C for 30 sec] and final extension at 72°C for 1 min. The amplification and quality of the ChIPseq libraries were checked by running 10% of the samples in E-Gel™ Agarose Gels with SYBR™ Safe DNA Gel Stain (ThermoFisher Scientific, USA), and if necessary, samples were reamplified additional 4 cycles using the same thermocycler protocol described above. Then, the libraries were purified and size-selected using Agencourt AMPure XP beads (1.25x vol. to remove short fragments. The concentration of ChIP-DNA libraries was measured by Qubit-4 fluorometer (ThermoFisher, USA) and equal amounts of each sample were pooled and 50bp paired-end reads were sequenced on an Illumina 4000 platform by GENEWIZ technology (GENEWIZ, USA).

### RNA-sequencing

For RNA-seq library preparation, in vitro polarized human Th1 cells either not stimulated or stimulated with the different IL-6 variants at 37°C for 6 hours, total RNA was extracted and RNAseq libraries were prepared by Edinburg Sequencing Core facility.

### ChIP-seq data analysis

The quality of generated libraries was inspected using FastQC v0.11.8. All sequencing reads were aligned to human reference genome (GRCh37; hg19) using bowtie.v1.2.2^1^ with default parameters except “--chunkmbs 1000 -S -m 1”. The genome index was generated using “bowtie-build” using default parameters. The aligned reads were indexed using samtools v1.9^2^ for further processing. The genome-wide binding profile (i.e. Bigwig files) were generated by bamCoverage v3.2.0^3^ using default parameters except “--normalizeUsing BPM -- minMappingQuality 30 --ignoreDuplicates --extendReads 250 --blackListFileName hg19.blacklist.bed”. The binding profiles were visualized using IGV genome browser v2.5.0^4^. Binding peaks were called by “callpeaks” procedure from MACS2 v2.1.2^5^ using default parameters except “-f BAMPE – nomodel -t mutant -c input”. The identified peaks were further screened against “hg19 blacklisted” genomic regions^6^, mitochondrial DNA, and pseudochromosomes. The binding heatmap surrounding HyIL-6 bound regions was generated by ChAsE v1.0.11^7^. HyIL-6 bound regions were sorted by significance and annotated by “annotatePeaks” procedure from HOMER v4.10^8^ to obtain the nearest genes. Pathway analysis of the top 2000 annotated genes was performed by Metascape^9^ on all GO terms related to biological processes. The resulting pathways were sorted by significance and plotted by Datagraph v4.3. The average binding signal intensity for each peak was calculated by UCSC bigWigAverageOverBed v2 using default parameters. *De novo* Motif findings were performed in 200bp bound regions (n=500) using MEME Suite v5.0.2^10^ with default parameters except “-maxsize 10000000 -dna -mod zoops -nmotifs 10”. *De novo* motifs were compared against all JASPAR known motifs by TOMTOM^11^. Statistical analyses were performed using the indicated Two-tailed parametric and non-parametric tests as appropriate.

### RNA-seq data analysis

The quality of generated libraries was inspected using FastQC v0.11.8. The RNA expression level in each library was estimated by “rsem-calculate-expression” procedure in RSEM v1.3.1^12^ using default parameters except “--bowtie-n 1 –bowtie-m 100 –seed-length 28 -- paired-end”. The bowtie index required by RSEM software was generated by “rsem-prepare-reference” on all RefSeq genes, obtained from UCSC table browser on April 2017. EdgeR v3.24.0^13^ package was used to normalize gene expression among all libraries and identify differentially expressed genes among samples with following constraints: fold change ≥ 1.5, FDR ≤0.05 and RPKM > 4 in at least one of two compared samples. The volcano plot representation was used to depict the log fold change of gene expression (Hy-IL6 vs. unstimulated; n=3) as a function of significance. The scatter plot was used to show the expression of genes in HyIL-6 stimulated against unstimulated samples. The expression values were the average of (n=3) independents donors. Differentially expressed genes under HyIL-6 stimulations were probed for response by the three indicated mutants (i.e. Mut3, C7, A1). The expression values were normalized to HyIL-6 and plotted by PRISM v8.1.0.

### T cells population differentiation

Resting CD4^+^ T cells isolated as described above were activated under Th1, Th17 or Tregs polarizing conditions. Briefly, resting human CD4^+^ T cells freshly isolated were activated using ImmunoCult™ Human CD3/CD28 T Cell Activator (StemCell, Cat#10971) following manufacturer instructions for 3 days in the presence of the cytokines required for the different CD4^+^ T cells populations: Th1 (IL2 (20 ng/ml), anti-IL4 (10 ng/ml), IL12 (20 ng/ml)), Th17 (IL1β (10 ng/ml, R&D Systems, Cat#201-LB/CF), IL23 (10 ng/ml, R&D Systems, Cat#1290-IL), anti-IL4 (10 ng/ml), anti-IFNγ (10 ng/ml, BD Biosciences, Cat#554698)) or Tregs (IL2 (20 ng/ml), TGF-β (5 ng/ml, Peprotech, Cat#100-21), anti-IL4 (10 ng/ml), anti-IFNγ (10 ng/ml)) in the presence or absence of saturating concentrations of the different variants of IL6 described in this manuscript. After three days of priming, cells were expanded for another 5 days in the presence of IL2 (20 ng/ml). Th1 and Th17 cells were restimulated for 6 hours in the presence of PMA (100 ng/ml, Sigma, Cat#P8139), lonomycin (1μM, Sigma, I0634) and Brefeldin A (5 μg/ml, Sigma, B7651) before FACS analysis. In all cases cells were fixed with 2% formaldehyde and prepared to be analysed by FACS. Cells were then permeabilised with Saponin 2% in PBS for 20 minutes at room temperature and then stained in Saponin 2% in PBS with the appropriate antibodies: Th1 (anti-CD3-BV510 (1:100, Biolegend, Cat#300448), anti-CD4-PE (1:100, Biolegend, Cat#357404), anti-CD8-AF700 (1:100, Biolegend, Cat#300920), anti-IFNγ (1:100, Biolegend, Cat#502217)), Th17 (anti-CD3-BV510, anti-CD4-PE, anti-CD8-AF700, anti-IL17A-APC (1:100, Biolegend, Cat#512334)) and Tregs (anti-CD3-BV510, anti-CD4-PE, anti-CD8-AF700, anti-CD25-APC (1:100, Biolegend, Cat#302610), anti-FoxP3-AF488 (1:100, Biolegend, 320012)) and analysed in a CytoFLEX S (Beckman Coulter).

### Live-cell dual-color single-molecule imaging studies

Single molecule imaging experiments were carried out by total internal reflection fluorescence (TIRF) microscopy with an inverted microscope (Olympus IX71) equipped with a triple-line total internal reflection (TIR) illumination condenser (Olympus) and a back-illuminated electron multiplied (EM) CCD camera (iXon DU897D, 512 x 512 pixel, Andor Technology). A 150 x magnification objective with a numerical aperture of 1.45 (UAPO 150 3 /1.45 TIRFM, Olympus) was used for TIR illumination. All experiments were carried out at room temperature in medium without phenol red supplemented with an oxygen scavenger and a redox-active photoprotectant to minimize photobleaching (Vogelsang et al., 2008). For cell surface labeling of mXFP-gp130, antiGFP-NB^DY647^ and antiGFP-NB^RHO11^ were added to the medium at equal concentrations (2 nM) and incubated for at least 5 min. The nanobodies were kept in the bulk solution during the whole experiment in order to ensure high equilibrium binding to mXFP-gp130. Dimerization of mXFP-gp130 was probed before and after incubation with either 50 nM HY-IL6 or 100 nM of the IL-6 mutants (Mut3, C7, A1). Image stacks of 150 frames were recorded at 32 ms/frame. For simultaneous dual color acquisition, antiGFP-NB^RHO11^ was excited by a 561 nm diode-pumped solid-state laser at 0.95 mW (~32W/cm2) and antiGFP-NB^DY647^ by a 642 nm laser diode at 0.65 mW (~22W/cm2). Fluorescence was detected using a spectral image splitter (DualView, Optical Insight) with a 640 DCXR dichroic beam splitter (Chroma) in combination with the bandpass filter 585/40 (Semrock) for detection of RHO11 and 690/70 (Chroma) for detection of DY647 dividing each emission channel into 512×256 pixel. In order to probe the dimerization/ligand binding of/to endogenous gp130 presented on HeLa cells, each ligand was (Hyper IL-6, Mut3, A1 & C7) conjugated to Dy547 and Dy647, respectivley. Prior the experiment untranfected HeLa cells were incubated with 10 nM of both (Dy547 and Dy647) dye-conjugated ligands for 10 minutes at room temperature and dual color experiments have been performed like above.

Single molecule localization and single molecule tracking were carried out using the multiple-target tracing (MTT) algorithm (Serge et al., 2008) as described previously (You et al., 2016). Step-length histograms were obtained from single molecule trajectories and fitted by two fraction mixture model of Brownian diffusion. Average diffusion constants were determined from the slope (2-10 steps) of the mean square displacement versus time lapse diagrams. Immobile molecules were identified by the density-based spatial clustering of applications with noise (DBSCAN) algorithm as described recently (Roder et al., 2014). For comparing diffusion properties and for co-tracking analysis, immobile particles were excluded from the data set. Prior to co-localization analysis, imaging channels were aligned with sub-pixel precision by using a spatial transformation. To this end, a transformation matrix was calculated based on a calibration measurement with multicolor fluorescent beads (TetraSpeck microspheres 0.1 mm, Invitrogen) visible in both spectral channels (cp2tform of type ‘affine’, The MathWorks MATLAB 2009a).

Individual molecules detected in the both spectral channels were regarded as co-localized, if a particle was detected in both channels of a single frame within a distance threshold of 100 nm radius. For single molecule co-tracking analysis, the MTT algorithm was applied to this dataset of co-localized molecules to reconstruct co-locomotion trajectories (co-trajectories) from the identified population of co-localizations. For the co-tracking analysis, only trajectories with a minimum of 10 steps (~300 ms) were considered. The relative fraction of co-tracked molecules was determined with respect to the absolute number of trajectories and corrected for gp130 stochastically double-labeled with the same fluorophore species as follows:

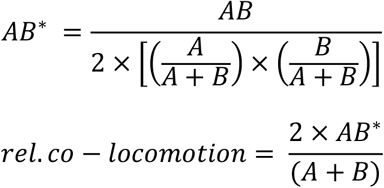

where A, B, AB and AB* are the numbers of trajectories observed for Rho11, DY647, co-trajectories and corrected co-trajectories, respectively.

### Imaging of receptors in endosomes

For tracing IL-6 uptake into early endosomes, HeLa cells were transiently transfected (PEI) with XFP-gp130 and seeded on 12mm cover slides placed in a 24-well plate. Cells were stimulated for 45 min at 37C with 20 nM of HY-IL6^DY547^ or 40 nM of Mut3^DY547^, C7^DY547^ and A1^DY547^ respectively. Cells were PFA fixed (4%,15 min in PBS), and washed 3x with PBS. Cells were permeabilized in a Methanol buffer (90% MeOH, MES, 10 mM EDTA 100 μM, MgCl_2_ 100 μM) for 1 min and washed 3x with PBS. Subsequently, cells were incubated in blocking buffer (TBS + 1% BSA = TBSA) for 20 min. Cells were incubated with the primary antibody against EEA1 (mouse-anti-human, 1:200, eBioscience, #14-9114-80) in TBSA for 45 min and washed 3x with TBSA. Cells were incubated with the secondary antibody (donkey-anti-mouse^Alexa647^ 1:200, Life Technologies, #A31571) in TBSA for 45 min, washed 3x with TBSA and mounted under coverslips using Vectashield mounting medium containing DAPI (Vector Laboratories) and viewed using an LSM 700 confocal microscope (Carl Zeiss).

## SUPPLEMENTARY FIGURE LEGENDS

**Supplementary Figure 1:**
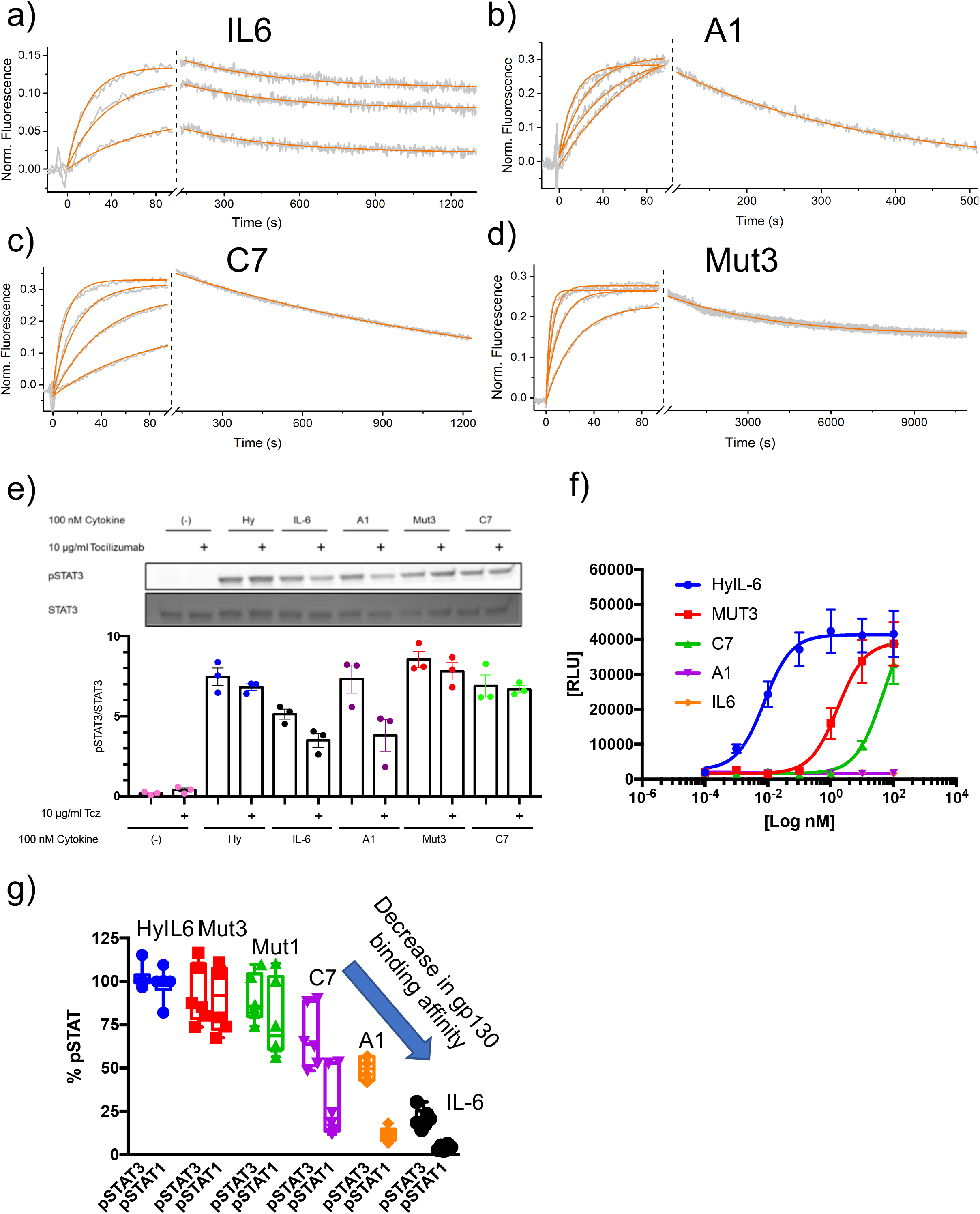
Biophysical characterization of IL-6 variants. (a-d) Switchsense binding sensograms for the different IL-6 ligands. Gp130 was loaded onto the chip and different concentrations of the indicated IL-6 ligands were injected. The binding constant values were estimated by fitting a pseudo first-order kinetic model. (e) HepG2 cells were pretreated with 10 μg/ml Tocilizumab (Tcz) as indicated for 30 min and then stimulated with 100 nM of the different IL-6 variants for 15 min. Phosphorylation of STAT3 was detected by Western Blot. One representative Western Blot and the quantification of three experiments is shown. (f) Ba/F3-gp130 cells were cultured in the presence of different concentrations of the IL-6 variants für 48h. Cell viability was assessed and is shown as relative light units (RLU). One representative experiment out of three with similar outcome is shown. (g) STAT1 and STAT3 activation levels induced by the indicated ligands in HeLa cells after 15 min stimulation measured by Phospho-Flow cytometry.

**Supplementary Figure 2:**
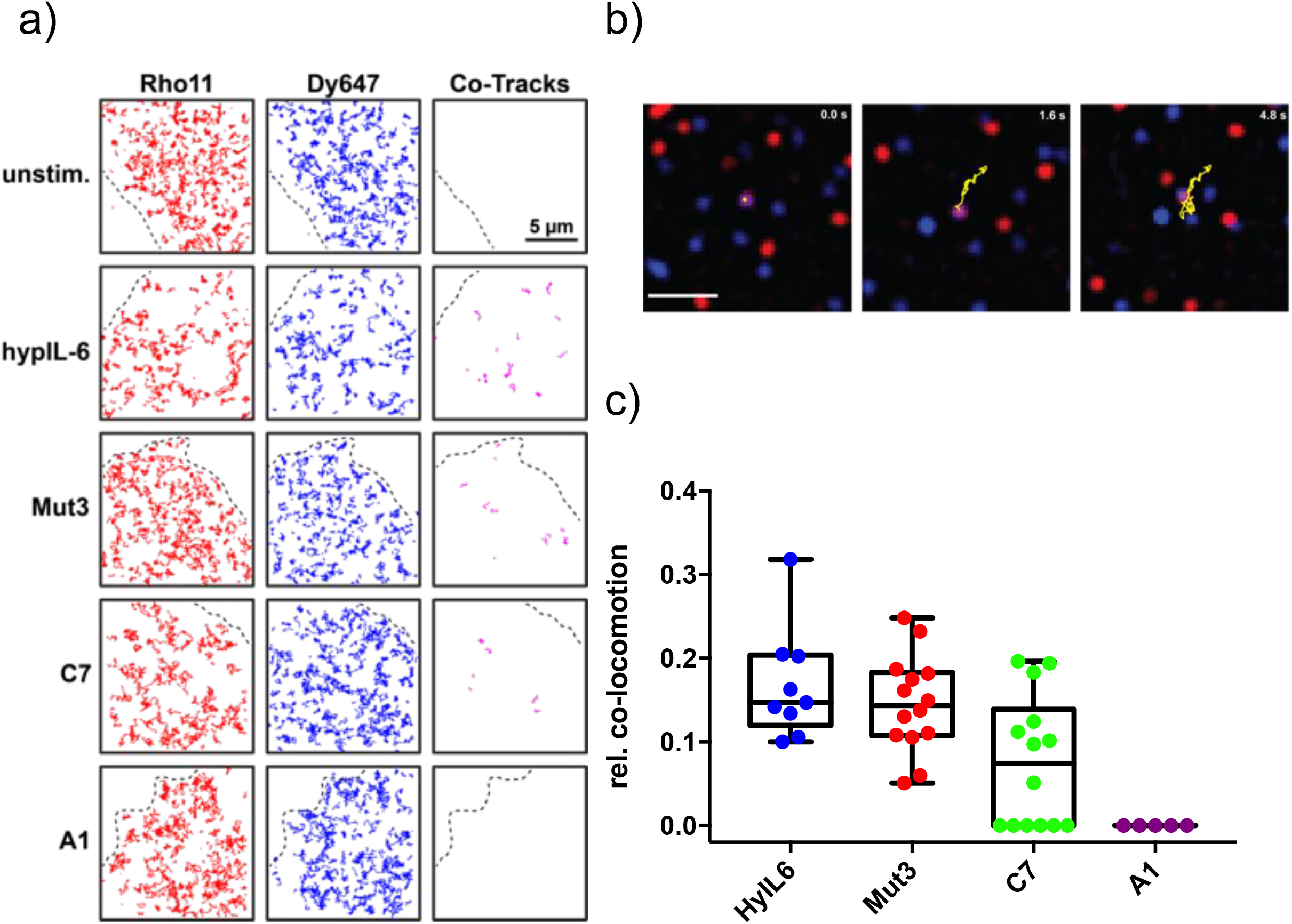
Dimerization of gp130 induced by IL-6 variants. (a)Trajectories of representative HEK 293T gp130KO cells, transiently expressing mXFP-gp130. Shown are the trajectories of the Rho11 (red), Dy647 (blue) and co-trajectories (magenta) for each condition. (b) Representative track after Mut3 stimulation over the entire 150 frames. (c) Co-locomotion analysis with dye-conjugated ligands bound to endogenous gp130 in HeLa cells (20 nM total ligand concentration).

**Supplementary Figure 3:**
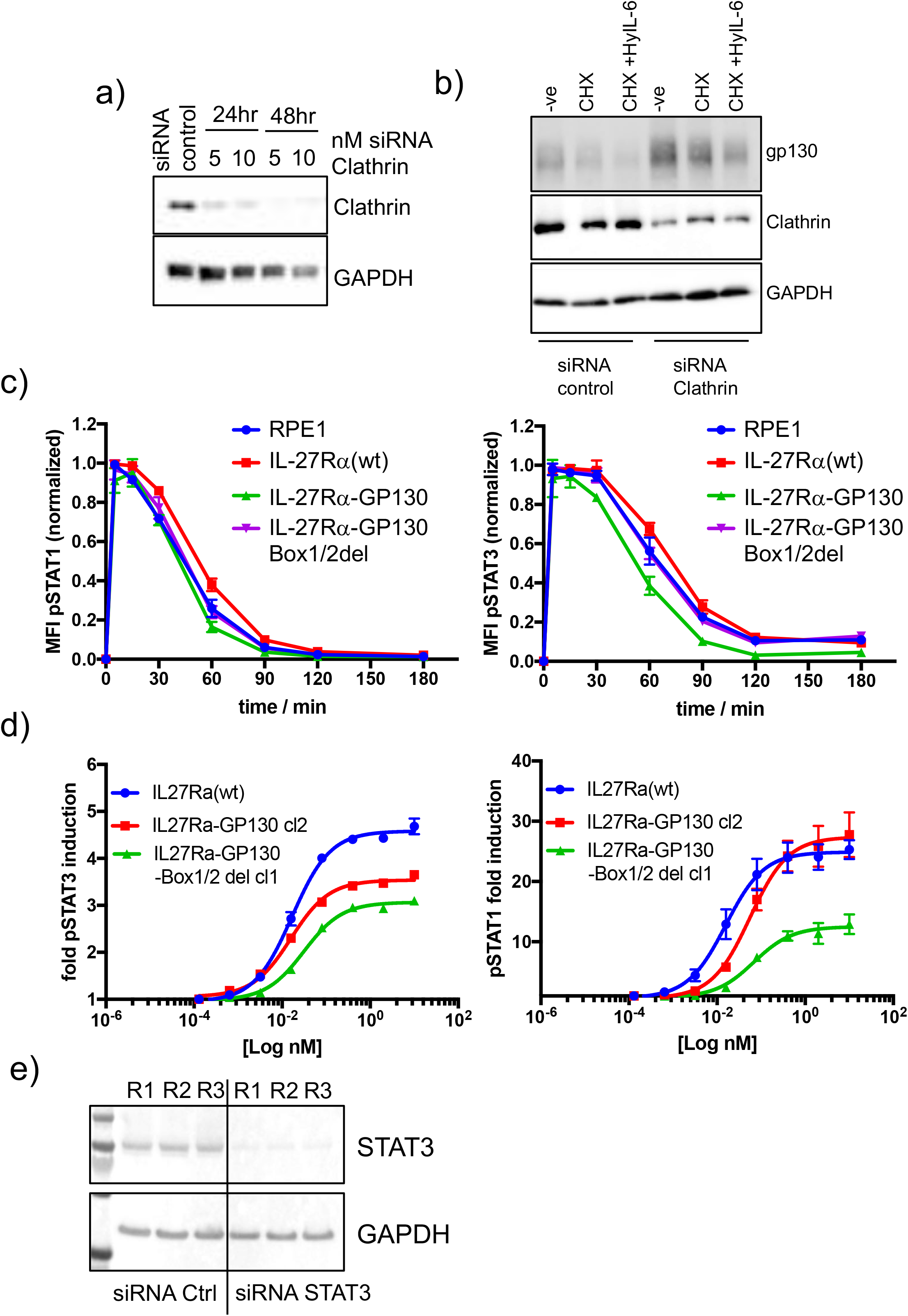
Functional characterization of RPE1 stable clones. (a) HeLa cells were transfected with different concentrations of siRNA targeting clathrin for 24 or 48 hours. Levels of clathrin were measured by western blot using clathrin specific antibodies. GAPDH was used as loading control. (b) HeLa cells were transfected with 10 nM control siRNA or clathrin specific siRNA. 48 hours later they were stimulated with saturating concentrations of HyIL-6 in the presence of cycloheximide for three hours. Gp130 and clathrin levels were measured by western blot. GAPDH was used as loading control. (c) The different RPE1 stable clones were stimulated with saturating concentrations of HyIL-6 for the indicated times and the levels of STAT1 (left panel) or STAT3 (right panel) activation were measured by phospho-Flow cytometry. Data are mean +/− SEM from three independent replicates, each perform in duplicate. (d) The different RPE1 stable clones were stimulated with the indicated doses of IL-27 for 15 min and the levels of STAT1 (left panel) and STAT3 (right panel) activation were measured by Phospho-Flow cytometry. Data are mean +/− SEM from three independent replicates, each perform in duplicate. (e) HeLa cells were transfected with control siRNA or STAT3 specific siRNA and 48 hours later levels of STAT3 were measured by western blot. GAPDH was used as loading control.

**Supplementary Figure 4:**
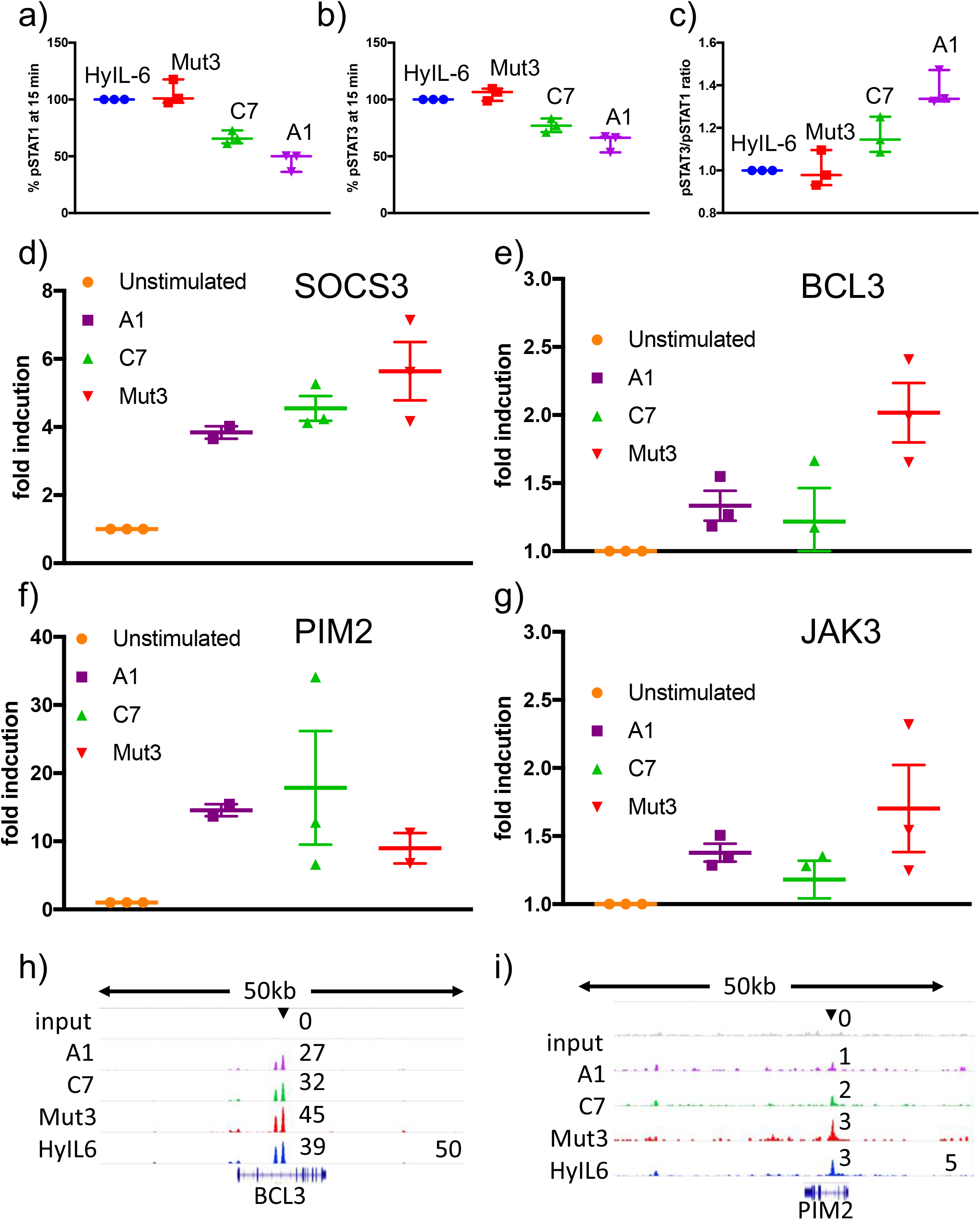
Transcriptional characterization of IL-6 variants. (a-c) Human CD4 T cells were isolated from buffy coats and differentiated into Th-1 cells for five days. 24 hr before stimulation, cells were starved of cytokines. Starved-Th-1 cells were stimulated with saturating doses of the indicated IL-6 ligads for 15 min and the levels of STAT1 (a) and STAT3 (b) activation were measured by Phospho-Flow cytometry. The pSTAT3/pSTAT1 ratio resulting from these studies is plotted in panel (c). Data are mean +/− SEM from three independent donors. (d-g) Th-1 cells generated as in (a-c) were stimulated with saturating concentrations of the indicated IL-6 ligands for six hours. RNA was extracted at that point and converted to cDNA to perform qPCRs studies. The levels of the indicated STAT3-induced genes were quantified. (h-i) STAT3 binding to *BCL3* (h) and *PIM2* (i) promoters in response to stimulation with the different IL-6 variants in Th-1 cells.

**Supplementary Figure 5:**
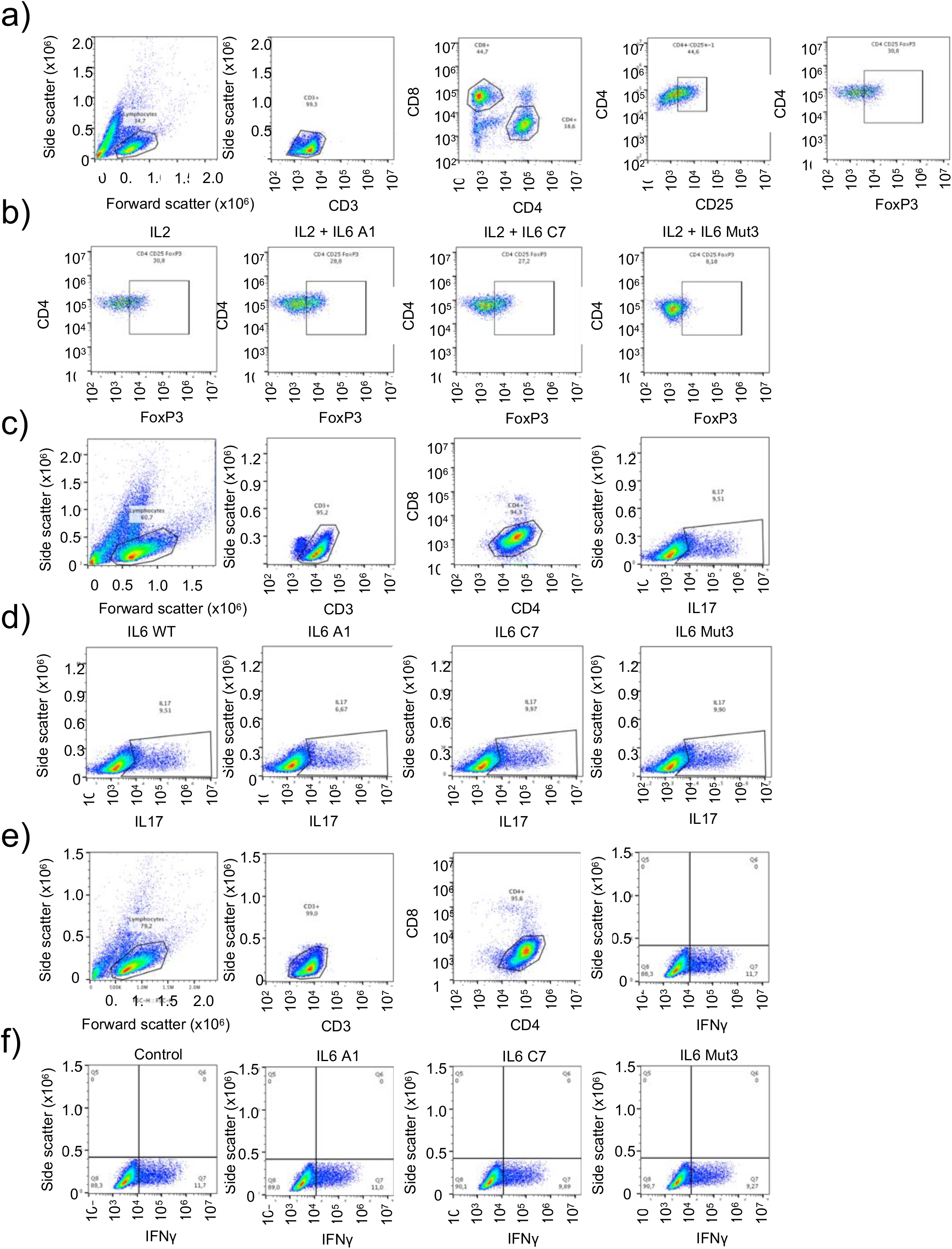
Immuno-modulatory properties of the IL-6 variants. (a-b) Representative FACS plots and population strategy to define Treg cell numbers in response to IL-6 variants. (c-d) Representative FACS plots and population strategy to define Th-17 cell numbers in response to IL-6 variants. (e-f) Representative FACS plots and population strategy to define Th-1 cell numbers in response to IL-6 variants.

**Supplementary Table 1.**
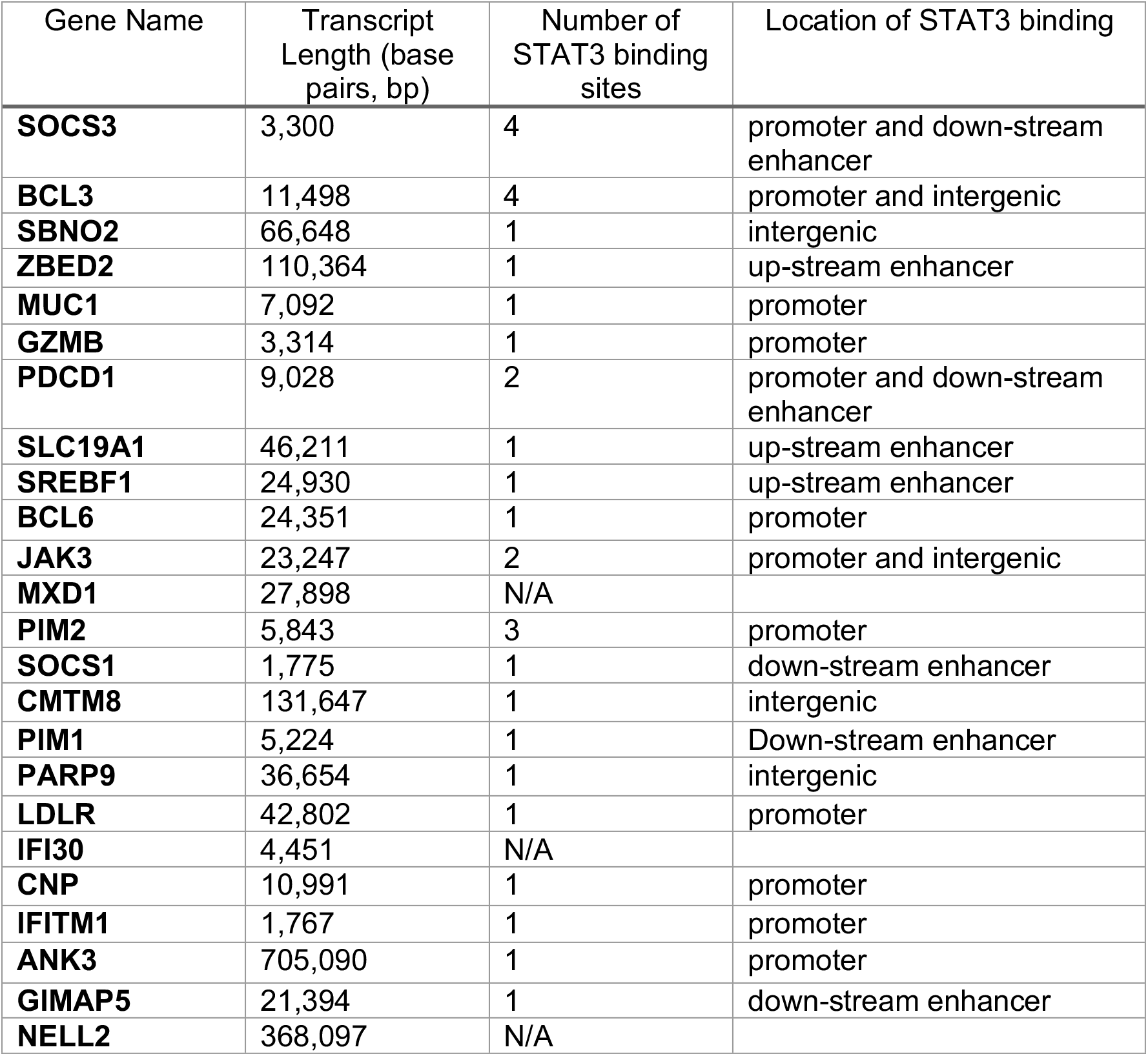

## REFERENCES

Acuto, O., Di Bartolo, V., and Michel, F. (2008). Tailoring T-cell receptor signals by proximal negative feedback mechanisms. Nat Rev Immunol 8, 699–712.

Becker, V., Schilling, M., Bachmann, J., Baumann, U., Raue, A., Maiwald, T., Timmer, J., and Klingmuller, U. Covering a broad dynamic range: information processing at the erythropoietin receptor. Science 328, 1404–1408.

Becker, V., Schilling, M., Bachmann, J., Baumann, U., Raue, A., Maiwald, T., Timmer, J., and Klingmuller, U. (2010). Covering a broad dynamic range: information processing at the erythropoietin receptor. Science 328, 1404–1408.

Boder, E.T., and Wittrup, K.D. (1997). Yeast surface display for screening combinatorial polypeptide libraries. Nat Biotechnol 15, 553–557.

Bonham, A.J., Wenta, N., Osslund, L.M., Prussin, A.J., 2nd, Vinkemeier, U., and Reich, N.O. (2013). STAT1:DNA sequence-dependent binding modulation by phosphorylation, protein:protein interactions and small-molecule inhibition. Nucleic Acids Res 41, 754–763.

Boulanger, M.J., Chow, D., Brevnova, E., Martick, M., Sandford, G., Nicholas, J., and Garcia, K.C. (2004). Molecular Mechanisms for Viral Mimicry of a Human Cytokine: Activation of gp130 by HHV-8 Interleukin-6. J Mol Biol 335, 641–654.

Bulut, G.B., Sulahian, R., Ma, Y., Chi, N.W., and Huang, L.J. Ubiquitination regulates the internalization, endolysosomal sorting, and signaling of the erythropoietin receptor. J Biol Chem 286, 6449–6457.

Cendrowski, J., Maminska, A., and Miaczynska, M. (2016). Endocytic regulation of cytokine receptor signaling. Cytokine Growth Factor Rev 32, 63–73.

Cheng, G., Yu, A., and Malek, T.R. (2011). T-cell tolerance and the multi-functional role of IL-2R signaling in T-regulatory cells. Immunol Rev 241, 63–76.

Claudinon, J., Monier, M.N., and Lamaze, C. (2007). Interfering with interferon receptor sorting and trafficking: impact on signaling. Biochimie 89, 735–743.

Costa-Pereira, A.P., Tininini, S., Strobl, B., Alonzi, T., Schlaak, J.F., Is’harc, H., Gesualdo, I., Newman, S.J., Kerr, I.M., and Poli, V. (2002). Mutational switch of an IL-6 response to an interferon-gamma-like response. Proc Natl Acad Sci U S A 99, 8043–8047.

Diehl, S., and Rincon, M. (2002). The two faces of IL-6 on Th1/Th2 differentiation. Mol Immunol 39, 531–536.

Ehret, G.B., Reichenbach, P., Schindler, U., Horvath, C.M., Fritz, S., Nabholz, M., and Bucher, P. (2001). DNA binding specificity of different STAT proteins. Comparison of in vitro specificity with natural target sites. J Biol Chem 276, 6675–6688.

Fallon, E.M., and Lauffenburger, D.A. (2000). Computational model for effects of ligand/receptor binding properties on interleukin-2 trafficking dynamics and T cell proliferation response. Biotechnol Prog 16, 905–916.

Fischer, M., Goldschmitt, J., Peschel, C., Brakenhoff, J.P., Kallen, K.J., Wollmer, A., Grotzinger, J., and Rose-John, S. (1997). I. A bioactive designer cytokine for human hematopoietic progenitor cell expansion. Nat Biotechnol 15, 142–145.

Gandhi, H., Worch, R., Kurgonaite, K., Hintersteiner, M., Schwille, P., Bokel, C., and Weidemann, T. (2014). Dynamics and interaction of interleukin-4 receptor subunits in living cells. Biophys J 107, 2515–2527.

Garbers, C., Thaiss, W., Jones, G.W., Waetzig, G.H., Lorenzen, I., Guilhot, F., Lissilaa, R., Ferlin, W.G., Grotzinger, J., Jones, S.A., et al. (2011). Inhibition of classic signaling is a novel function of soluble glycoprotein 130 (sgp130), which is controlled by the ratio of interleukin 6 and soluble interleukin 6 receptor. J Biol Chem 286, 42959–42970.

Gearing, D.P., Ziegler, S.F., Comeau, M.R., Friend, D., Thoma, B., Cosman, D., Park, L., and Mosley, B. (1994). Proliferative responses and binding properties of hematopoietic cells transfected with low-affinity receptors for leukemia inhibitory factor, oncostatin M, and ciliary neurotrophic factor. Proc Natl Acad Sci U S A 91, 1119–1123.

German, C.L., Sauer, B.M., and Howe, C.L. The STAT3 beacon: IL-6 recurrently activates STAT 3 from endosomal structures. Exp Cell Res 317, 1955–1969.

Gil, M.P., Bohn, E., O’Guin, A.K., Ramana, C.V., Levine, B., Stark, G.R., Virgin, H.W., and Schreiber, R.D. (2001). Biologic consequences of Stat1-independent IFN signaling. Proc Natl Acad Sci U S A 98, 6680–6685.

Gonnord, P., Blouin, C.M., and Lamaze, C. (2012). Membrane trafficking and signaling: two sides of the same coin. Semin Cell Dev Biol 23, 154–164.

Gorby, C., Martinez-Fabregas, J., Wilmes, S., and Moraga, I. (2018). Mapping Determinants of Cytokine Signaling via Protein Engineering. Front Immunol 9, 2143.

Grotzinger, J., Kurapkat, G., Wollmer, A., Kalai, M., and Rose-John, S. (1997). The family of the IL-6-type cytokines: specificity and promiscuity of the receptor complexes. Proteins 27, 96–109.

Heink, S., Yogev, N., Garbers, C., Herwerth, M., Aly, L., Gasperi, C., Husterer, V., Croxford, A.L., Moller-Hackbarth, K., Bartsch, H.S., et al. (2017). Trans-presentation of IL-6 by dendritic cells is required for the priming of pathogenic TH17 cells. Nat Immunol 18, 74–85.

Heller, N.M., Qi, X., Junttila, I.S., Shirey, K.A., Vogel, S.N., Paul, W.E., and Keegan, A.D. (2008). Type I IL-4Rs selectively activate IRS-2 to induce target gene expression in macrophages. Sci Signal 1, ra17.

Herrero, C., Hu, X., Li, W.P., Samuels, S., Sharif, M.N., Kotenko, S., and Ivashkiv, L.B. (2003). Reprogramming of IL-10 activity and signaling by IFN-gamma. J Immunol 171, 5034–5041.

Hilger, D., Masureel, M., and Kobilka, B.K. (2018). Structure and dynamics of GPCR signaling complexes. Nat Struct Mol Biol 25, 4–12.

Ho, C.C.M., Chhabra, A., Starkl, P., Schnorr, P.J., Wilmes, S., Moraga, I., Kwon, H.S., Gaudenzio, N., Sibilano, R., Wehrman, T.S., et al. (2017). Decoupling the Functional Pleiotropy of Stem Cell Factor by Tuning c-Kit Signaling. Cell 168, 1041–1052 e1018.

Horvath, C.M., Wen, Z., and Darnell, J.E., Jr. (1995). A STAT protein domain that determines DNA sequence recognition suggests a novel DNA-binding domain. Genes Dev 9, 984–994.

Hunter, C.A., and Jones, S.A. (2015). IL-6 as a keystone cytokine in health and disease. Nat Immunol 16, 448–457.

Jones, G.W., McLoughlin, R.M., Hammond, V.J., Parker, C.R., Williams, J.D., Malhotra, R., Scheller, J., Williams, A.S., Rose-John, S., Topley, N., et al. (2010). Loss of CD4+ T cell IL-6R expression during inflammation underlines a role for IL-6 trans signaling in the local maintenance of Th17 cells. J Immunol 184, 2130–2139.

Keeler, C., Jablonski, E.M., Albert, Y.B., Taylor, B.D., Myszka, D.G., Clevenger, C.V., and Hodsdon, M.E. (2007). The kinetics of binding human prolactin, but not growth hormone, to the prolactin receptor vary over a physiologic pH range. Biochemistry 46, 2398–2410.

Kim, A.R., Ulirsch, J.C., Wilmes, S., Unal, E., Moraga, I., Karakukcu, M., Yuan, D., Kazerounian, S., Abdulhay, N.J., King, D.S., et al. (2017). Functional Selectivity in Cytokine Signaling Revealed Through a Pathogenic EPO Mutation. Cell 168, 1053–1064 e1015.

Kimura, A., and Kishimoto, T. (2010). IL-6: regulator of Treg/Th17 balance. Eur J Immunol 40, 1830–1835.

Korn, T., Mitsdoerffer, M., Croxford, A.L., Awasthi, A., Dardalhon, V.A., Galileos, G., Vollmar, P., Stritesky, G.L., Kaplan, M.H., Waisman, A., et al. (2008). IL-6 controls Th17 immunity in vivo by inhibiting the conversion of conventional T cells into Foxp3+ regulatory T cells. Proc Natl Acad Sci U S A 105, 18460–18465.

Krutzik, P.O., and Nolan, G.P. (2006). Fluorescent cell barcoding in flow cytometry allows high-throughput drug screening and signaling profiling. Nat Methods 3, 361–368.

LaPorte, S.L., Juo, Z.S., Vaclavikova, J., Colf, L.A., Qi, X., Heller, N.M., Keegan, A.D., and Garcia, K.C. (2008). Molecular and structural basis of cytokine receptor pleiotropy in the interleukin-4/13 system. Cell 132, 259–272.

Lemmon, M.A., and Schlessinger, J. (2010). Cell signaling by receptor tyrosine kinases. Cell 141, 1117–1134.

Louten, J., Boniface, K., and de Waal Malefyt, R. (2009). Development and function of TH17 cells in health and disease. J Allergy Clin Immunol 123, 1004–1011.

Marchetti, M., Monier, M.N., Fradagrada, A., Mitchell, K., Baychelier, F., Eid, P., Johannes, L., and Lamaze, C. (2006). Stat-mediated signaling induced by type I and type II interferons (IFNs) is differentially controlled through lipid microdomain association and clathrin-dependent endocytosis of IFN receptors. Mol Biol Cell 17, 2896–2909.

McKeithan, T.W. (1995). Kinetic proofreading in T-cell receptor signal transduction. Proc Natl Acad Sci U S A 92, 5042–5046.

Mendoza, J.L., Schneider, W.M., Hoffmann, H.H., Vercauteren, K., Jude, K.M., Xiong, A., Moraga, I., Horton, T.M., Glenn, J.S., de Jong, Y.P., et al. (2017). The IFN-lambda-IFN-lambdaR1-IL-10Rbeta Complex Reveals Structural Features Underlying Type III IFN Functional Plasticity. Immunity 46, 379–392.

Moraga, I., Richter, D., Wilmes, S., Winkelmann, H., Jude, K., Thomas, C., Suhoski, M.M., Engleman, E.G., Piehler, J., and Garcia, K.C. (2015a). Instructive roles for cytokine-receptor binding parameters in determining signaling and functional potency. Sci Signal 8, ra114.

Moraga, I., Wernig, G., Wilmes, S., Gryshkova, V., Richter, C.P., Hong, W.J., Sinha, R., Guo, F., Fabionar, H., Wehrman, T.S., et al. (2015b). Tuning cytokine receptor signaling by reorienting dimer geometry with surrogate ligands. Cell 160, 1196–1208.

Murray, P.J. (2007). The JAK-STAT signaling pathway: input and output integration. J Immunol 178, 2623–2629.

Naka, T., Nishimoto, N., and Kishimoto, T. (2002). The paradigm of IL-6: from basic science to medicine. Arthritis Res 4 Suppl 3, S233–242.

Payelle-Brogard, B., and Pellegrini, S. Biochemical monitoring of the early endocytic traffic of the type I interferon receptor. J Interferon Cytokine Res 30, 89–98.

Pestka, S. (2007). The interferons: 50 years after their discovery, there is much more to learn. The Journal of biological chemistry 282, 20047–20051.

Piehler, J., Thomas, C., Garcia, K.C., and Schreiber, G. (2012). Structural and dynamic determinants of type I interferon receptor assembly and their functional interpretation. Immunol Rev 250, 317–334.

Reddy, C.C., Niyogi, S.K., Wells, A., Wiley, H.S., and Lauffenburger, D.A. (1996). Engineering epidermal growth factor for enhanced mitogenic potency. Nat Biotechnol 14, 1696–1699.

Ring, A.M., Lin, J.X., Feng, D., Mitra, S., Rickert, M., Bowman, G.R., Pande, V.S., Li, P., Moraga, I., Spolski, R., et al. (2012). Mechanistic and structural insight into the functional dichotomy between IL-2 and IL-15. Nat Immunol 13, 1187–1195.

Rochman, Y., Spolski, R., and Leonard, W.J. (2009). New insights into the regulation of T cells by gamma(c) family cytokines. Nat Rev Immunol 9, 480–490.

Roder, F., Lubk, A., Wolf, D., and Niermann, T. (2014). Noise estimation for off-axis electron holography. Ultramicroscopy 144, 32–42.

Sarkar, C.A., Lowenhaupt, K., Horan, T., Boone, T.C., Tidor, B., and Lauffenburger, D.A. (2002). Rational cytokine design for increased lifetime and enhanced potency using pH-activated “histidine switching”. Nat Biotechnol 20, 908–913.

Schindler, C., Levy, D.E., and Decker, T. (2007). JAK-STAT signaling: from interferons to cytokines. J Biol Chem 282, 20059–20063.

Schmitz, J., Dahmen, H., Grimm, C., Gendo, C., Muller-Newen, G., Heinrich, P.C., and Schaper, F. (2000). The cytoplasmic tyrosine motifs in full-length glycoprotein 130 have different roles in IL-6 signal transduction. J Immunol 164, 848–854.

Schwerd, T., Twigg, S.R.F., Aschenbrenner, D., Manrique, S., Miller, K.A., Taylor, I.B., Capitani, M., McGowan, S.J., Sweeney, E., Weber, A., et al. (2017). A biallelic mutation in IL6ST encoding the GP130 co-receptor causes immunodeficiency and craniosynostosis. J Exp Med 214, 2547–2562.

Serge, A., Bertaux, N., Rigneault, H., and Marguet, D. (2008). Dynamic multiple-target tracing to probe spatiotemporal cartography of cell membranes. Nat Methods 5, 687–694.

Shah, M., Patel, K., Mukhopadhyay, S., Xu, F., Guo, G., and Sehgal, P.B. (2006). Membrane-associated STAT3 and PY-STAT3 in the cytoplasm. J Biol Chem 281, 7302–7308.

Spangler, J.B., Moraga, I., Mendoza, J.L., and Garcia, K.C. (2015). Insights into cytokine-receptor interactions from cytokine engineering. Annu Rev Immunol 33, 139–167.

Stroud, R.M., and Wells, J.A. (2004). Mechanistic diversity of cytokine receptor signaling across cell membranes. Sci STKE 2004, re7.

Stumhofer, J.S., Tait, E.D., Quinn, W.J., 3rd, Hosken, N., Spudy, B., Goenka, R., Fielding, C.A., O’Hara, A.C., Chen, Y., Jones, M.L., et al. A role for IL-27p28 as an antagonist of gp130-mediated signaling. Nat Immunol 11, 1119–1126.

Subramaniam, P.S., Khan, S.A., Pontzer, C.H., and Johnson, H.M. (1995). Differential recognition of the type I interferon receptor by interferons tau and alpha is responsible for their disparate cytotoxicities. Proc Natl Acad Sci U S A 92, 12270–12274.

Tanaka, Y., Tanaka, N., Saeki, Y., Tanaka, K., Murakami, M., Hirano, T., Ishii, N., and Sugamura, K. (2008). c-Cbl-dependent monoubiquitination and lysosomal degradation of gp130. Mol Cell Biol 28, 4805–4818.

Villasenor, R., Kalaidzidis, Y., and Zerial, M. (2016). Signal processing by the endosomal system. Curr Opin Cell Biol 39, 53–60.

Vogelsang, J., Kasper, R., Steinhauer, C., Person, B., Heilemann, M., Sauer, M., and Tinnefeld, P. (2008). A reducing and oxidizing system minimizes photobleaching and blinking of fluorescent dyes. Angew Chem Int Ed Engl 47, 5465–5469.

Waichman, S., Bhagawati, M., Podoplelova, Y., Reichel, A., Brunk, A., Paterok, D., and Piehler, J. (2010). Functional immobilization and patterning of proteins by an enzymatic transfer reaction. Anal Chem 82, 1478–1485.

Waldmann, T.A. (2006). The biology of interleukin-2 and interleukin-15: implications for cancer therapy and vaccine design. Nat Rev Immunol 6, 595–601.

Walter, M.R. (2004). Structural analysis of IL-10 and Type I interferon family members and their complexes with receptor. Advances in protein chemistry 68, 171–223.

Wang, X., Lupardus, P., Laporte, S.L., and Garcia, K.C. (2009). Structural biology of shared cytokine receptors. Annu Rev Immunol 27, 29–60.

Wells, J.A., Cunningham, B.C., Fuh, G., Lowman, H.B., Bass, S.H., Mulkerrin, M.G., Ultsch, M., and deVos, A.M. (1993). The molecular basis for growth hormone-receptor interactions. Recent Prog Horm Res 48, 253–275.

Wiederkehr-Adam, M., Ernst, P., Muller, K., Bieck, E., Gombert, F.O., Ottl, J., Graff, P., Grossmuller, F., and Heim, M.H. (2003). Characterization of phosphopeptide motifs specific for the Src homology 2 domains of signal transducer and activator of transcription 1 (STAT1) and STAT3. J Biol Chem 278, 16117–16128.

Wilmes, S., Beutel, O., Li, Z., Francois-Newton, V., Richter, C.P., Janning, D., Kroll, C., Hanhart, P., Hotte, K., You, C., et al. (2015). Receptor dimerization dynamics as a regulatory valve for plasticity of type I interferon signaling. J Cell Biol 209, 579–593.

Wilson, E.H., Wille-Reece, U., Dzierszinski, F., and Hunter, C.A. (2005). A critical role for IL-10 in limiting inflammation during toxoplasmic encephalitis. J Neuroimmunol 165, 63–74.

Wootten, D., Christopoulos, A., Marti-Solano, M., Babu, M.M., and Sexton, P.M. (2018). Mechanisms of signalling and biased agonism in G protein-coupled receptors. Nat Rev Mol Cell Biol 19, 638–653.

Wung, B.S., Ni, C.W., and Wang, D.L. (2005). ICAM-1 induction by TNFalpha and IL-6 is mediated by distinct pathways via Rac in endothelial cells. J Biomed Sci 12, 91–101.

You, C., Marquez-Lago, T.T., Richter, C.P., Wilmes, S., Moraga, I., Garcia, K.C., Leier, A., and Piehler, J. (2016). Receptor dimer stabilization by hierarchical plasma membrane microcompartments regulates cytokine signaling. Sci Adv 2, e1600452.

Zhao, W., Lee, C., Piganis, R., Plumlee, C., de Weerd, N., Hertzog, P.J., and Schindler, C. (2008). A conserved IFN-alpha receptor tyrosine motif directs the biological response to type I IFNs. J Immunol 180, 5483–5489.

Zinkle, A., and Mohammadi, M. (2018). A threshold model for receptor tyrosine kinase signaling specificity and cell fate determination. F1000Res 7.

